# 3D Bioprinting of Human Neural Tissues with Functional Connectivity

**DOI:** 10.1101/2024.01.18.576289

**Authors:** Yuanwei Yan, Xueyan Li, Yu Gao, Sakthikumar Mathivanan, Linghai Kong, Yunlong Tao, Yi Dong, Xiang Li, Anita Bhattacharyya, Xinyu Zhao, Su-Chun Zhang

## Abstract

Probing how the human neural networks operate is hindered by the lack of reliable human neural tissues amenable for dynamic functional assessment of neural circuits. We developed a 3D bioprinting platform to assemble tissues with defined human neural cell types in a desired dimension using a commercial bioprinter. The printed neuronal progenitors differentiate to neurons and form functional neural circuits in and between tissue layers with specificity within weeks, evidenced by the cortical-to-striatal projection, spontaneous synaptic currents and synaptic response to neuronal excitation. Printed astrocyte progenitors develop into mature astrocytes with elaborated processes and form functional neuron-astrocyte networks, indicated by calcium flux and glutamate uptake in response to neuronal excitation under physiological and pathological conditions. These designed human neural tissues will likely be useful for understanding the wiring of human neural networks, modeling pathological processes, and serving as platforms for drug testing.

## INTRODUCTION

The human brain forms by the integration of specialized neuronal and glial types that are precisely wired into networks. Understanding how the human neural networks operate is essential to probing the health and disease of our brain. Animal models cannot precisely recapitulate the high-order information processing of the human brain due to the differences in neuronal composition, synaptic integration, astrocytic complexity and neural networks between animals and humans (Beaulieu-Laroche et al., 2018; Sousa et al., 2017; Vasile et al., 2017). There is therefore a need of a reliable model of living human neural tissues amenable for functional network assessment.

Human neural tissues with 3D cytoarchitectures have been engineered from human pluripotent stem cells (hPSCs), including induced pluripotent stem cells (iPSCs) and embryonic stem cells (hESCs), using culture systems like hydrogel cultures (Choi et al., 2014; Madl et al., 2017), scaffold-based cultures (Tang-Schomer et al., 2014; Yan et al., 2017), acoustic levitational assembly (Bouyer et al., 2016), self-organized brain spheroid/organoid (Lancaster et al., 2013; Pasca et al., 2015), and brain-on-a-chip models (Harberts et al., 2020; Vatine et al., 2019). The 3D bioprinting represents more precisely controlled and defined technology for fabricating human neural tissues by spatial deposition of living cells and hydrogels into a biologically complex cytoarchitecture (Murphy and Atala, 2014; Qiu et al., 2020). However, printing soft tissues like the brain structure is difficult due to the inability of soft biomaterials to support a complex 3D structure or stiff gels to enable function. Although a scaffold provides a supporting structure for printing neural tissues (Crowe et al., 2020; Koroleva et al., 2021; Pagan-Diaz et al., 2019), it is not real 3D bioprinting or cellular bioprinting via direct deposition of live cells together with the bioink. In these cultures, the scaffold or mold is first fabricated by 3D pirnitng and then the cells are seeded on the scaffold or mold. Neural cells in the scaffold or mold were not evenly distributed and formed large and thick aggregates or cell clusters (Koroleva et al., 2021; Pagan-Diaz et al., 2019). More importantly, the scaffolds and molds are not biodegradable and block neural cell migration and in particular, neural network formation between layers. The 3D bioprinted neural tissues, fabricated using soft gel like gellan gum (Lozano et al., 2015), alginate (Joung et al., 2018), or gelatin mixed with fibrin (Noh et al., 2020), and printed layer-by-layer vertically, exhibit layered structures with certain degree of neuronal maturation. However, printed human neural tissues with functional neuron-neuron or neuron-glial connectivity within or between tissue layers were not shown (Table S1).

Here, we present a technology platform for assembling neuronal and glial subtypes into defined 3D neural tissues in which neurons and glia formed functional connections in and between tissue layers using extrusion bioprinting. This is achieved by printing one layer or band next to another horizontally rather than the traditional way of stacking the layers vertically. These specially designed 3D neural tissues can be maintained by the conventional culture systems and amenable for easy live-cell imaging and electrophysiological recording, providing a new platform for examining human neural networks under physiological and pathological conditions.

## DESIGN

Our goal is to construct a layered neural tissue in which neural progenitor cells mature and form synapses within and across layers while the structure is maintained. The biomaterial for tissue printing, or bioink, is the key for the success of bioprinting. The properties of this bioink need 1) to support the survival of printed cells, 2) to promote the functional maturation of neural cells, 3) to sustain designed cell distribution, 4) to possess adjustable gelation time, and 5) to maintain the structural stability of the gel. The fibrin hydrogel, consisting primarily of fibrinogen and thrombin, has been shown to be biocompatible for neural cells (Bouyer et al., 2016; Noh et al., 2020; Xu et al., 2006). Thus, we choose fibrin gel as the basic bioink in our printing process.

The bioprinting strateties include droplet-based, extrusion-based and lased-based methods. The extrusion 3D bioprinting deposits gel layer by layer and can be utilized to mimic the brain structure, like the laminations of the human cortex. Fabrication of multi-layered brain tissues is typically performed by stacking layers or assembling layer by layer vertically (Joung et al., 2018; Lee et al., 2009; Lozano et al., 2015). Construction of such kind of tissues requires the support with stiff gels which, however, inhibit the growth of neurites (nerves) and, in particular, the functional connections (e.g., synapses) between neural cells (Georges et al., 2006; Koser et al., 2016). The dimension of the printed tissue is another key factor in desiging the 3D bioprinting. The diffusion limit of oxygen is limited to roughly 100-200 μm for avascular tissues (Rademakers et al., 2019). The “thick” neural tissues limit the transfer of oxygen and nutrients, imparing the growth and function of the printed neural cells without the vascular system. Thus, the ideal thickness of the printed brain tissue would be around 100-200 μm. Further considerations in the design included the ease of morphological and functional assays as well as broad application in ordinary laboratories. We hence chose a thickness of 50 µm for each layer. In addition, unlike the traditional ways of stacking the tissue layers vertically (Lozano et al., 2015), we constructed a multi-layered tissue by depositing the layer horizontally next to each other. Together, this design enabled construction of a relatively thin but multi-layered and functional neural tissue with defined cellular compositions and desired dimensions, which can be maintained and assayed with ease in an ordinary laboratory.

## RESULTS

### Developing bioink for printing human neural tissues

We first identified an optimal concentration of fibrinogen and thrombin by measuring the survival of hPSC-derived cortical neural progenitor cells (NPCs) (Figure 1A-1C, and S1A-S1F). The cell viability decreased with the increasing thrombin concentrations at the fixed fibrinogen concentration of 2.5mg/mL, while it was not influenced by the increasing concentrations of fibrinogen at the fixed low concentration of thrombin (0.5U) (Figure S1E and S1F). The cells, however, tended to aggregate at higher fibrinogen levels (data not shown). We hence chose 2.5 or 5 mg/mL fibrinogen with 0.5 or 1 U thrombin for identifying an optimal gelation time.

**Figure 1.**
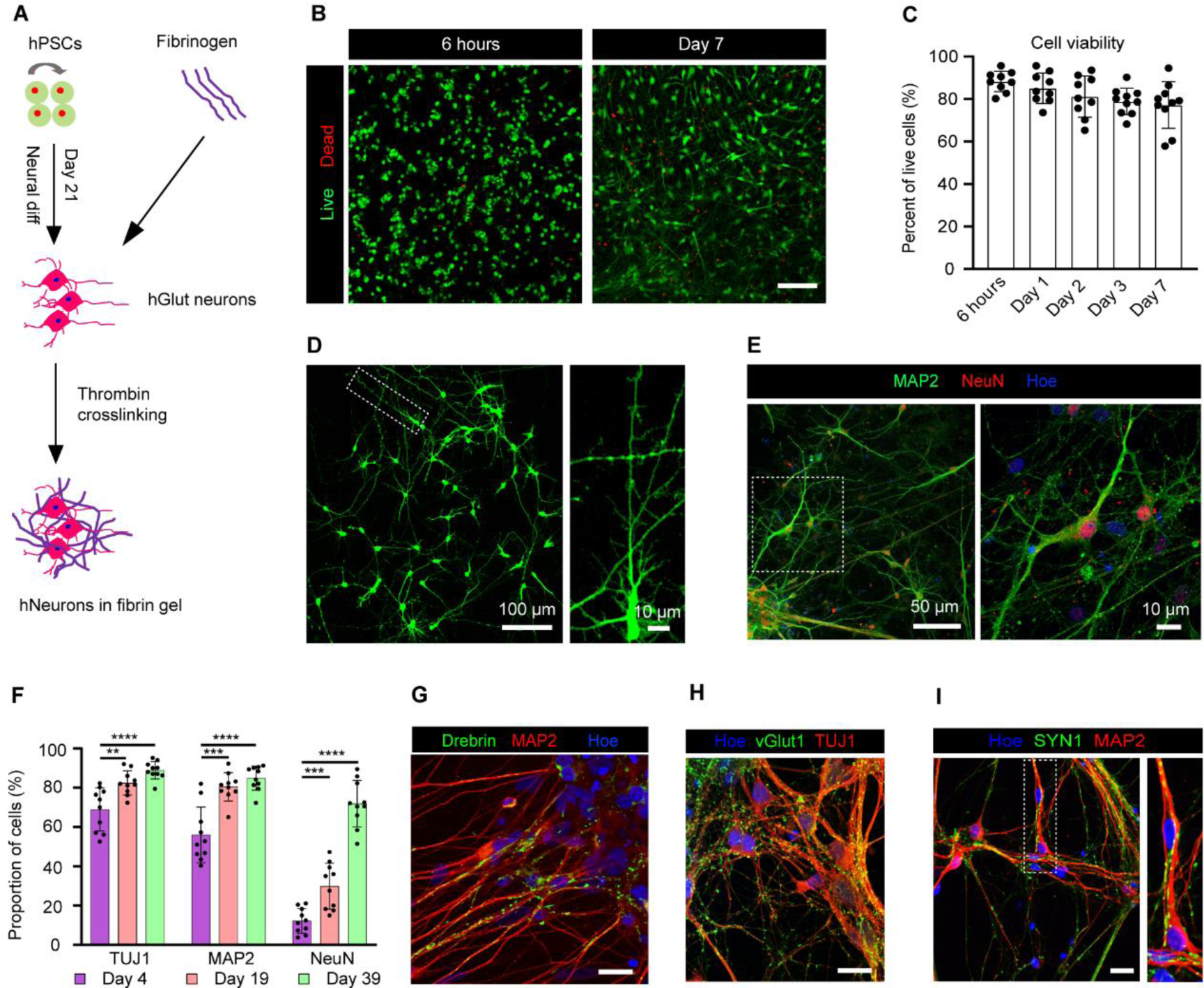
Survival and differentiation of hPSC-derived neurons in fibrin hydrogel. (A) Schematic diagram illustrating the growth of hPSC-derived NPCs in fibrin gel. hPSCs, human pluripotent stem cells; diff, differentiation; NPCs, neural progenitor cells; hGlut neurons, human cortical glutamatergic neurons. hNeurons, human neurons. (B) Live and Dead staining (green: “live”, red: “dead”) of neural cells grown in fibrin gel at 6 hours and day 7. (C) Viability (percentage of live cells) of neural cells at different time points (n= 9-10 from at least three different experiments). (D) Neurons showed a pyramidal morphology and formed dendritic spine-like structures, revealed by GFP after one month of growth in the gel. (E) Immunostaining for neuronal markers MAP2 and NeuN at day 39. (F) Quantitative analysis of the expression of TUJ1, MAP2 and NeuN for day 4, day 19 and 39 in the fibrin gel. (n=10 from at least three experiments. TUJ1, day 4 vs day 19, multiple t-test, ** P < 0.01; day 4 vs day 39, multiple t-test, **** P < 0.0001. MAP2, day 4 vs day 19, multiple t-test, *** P < 0.001; day 4 vs day 39, multiple t-test, **** P < 0.0001. NeuN, day 4 vs day 19, multiple t-test, *** P < 0.001; day 4 vs day 39, multiple t-test, **** P < 0.0001.) (G) Immunostaining for dendritic spine marker Drebrin with MAP2 at day 43 in the gel. (H) and (I) Immunostaining for synaptic markers vGlut1, SYN1 with TUJ1 or MAP2 at day 35 in the fibrin gel. Data are represented as mean ± SEM. Hoe: Hoechst 33342. Scale bars, 200 µm (B), 20 µm (G and H), 10 µm (I). See also Figure S1.

We then assessed the optimal gelation time. At a constant concentration of fibrinogen (5 or 2.5 mg/mL), the gelation time decreased with the increased level of thrombin (Figure S1C). At 0.5 U thrombin, the gelation time also reduced with increasing fibrinogen concentrations (Figure S1C). We chose 2.5 mg/mL fibrinogen and 0.5 U thrombin for hydrogel construction, yielding a gelation time of about 145 ± 10 seconds (Figure S1C), which would be sufficient for printing a 24-well plate. Under this condition, more than 85% of the cells survived after 6 hours of culturing in the fibrin gel and about 80% of the cells retained viable for 7 days (Figure 1B and 1C). We, therefore, used this composition of fibrin gel for the following experiments.

The fibrin gel itself has a low printability due to its high viscosity. We hence explored to combine with other hydrogels that are commonly used for printing (Caliari and Burdick, 2016; England et al., 2017; Holzl et al., 2016; Hospodiuk et al., 2017), including gelatin, alginate, Matrigel, hyaluronic acid, and nanofibrillated cellulose (Table S2). Matrigel and hyaluronic acid presented a high cell viability of 92.19 ± 4.26% and 85.13 ± 6.05%, respectively, although Matrigel frequently clogged the nozzle (Figure S1D, and Table S2). Thus, we developed a printable bioink based on the fibrin gel mixed with hyaluronic acid for our neural tissue printing.

A special requirement for neural tissue printing is that the bioink must support neurite growth and synaptogenesis. The cortical NPCs, differentiated from GFP- or mCherry-labeled hESCs for 21 days, expressed forebrain cortical progenitor markers FOXG1 and PAX6 (Figure S1A and S2B). When cultured in the fibrin gel, the progenitors became process-bearing (Figure 1D and S1G), TUJ1^+^ neurons (Figure S1H) as early as at day 4 after printing and progressively expressed mature neuronal markers MAP2 and NeuN (Figure 1E). Many neurons displayed a pyramidal morphology after 1 month of culture in the gel (Figure 1E and 1F). Importantly, the neurons extended elaborate neurites with fine dendritic structures, revealed by GFP labeling (Figure 1D). This was confirmed by positive staining for drebrin, a dendritic spine marker at day 40 (Figure 1G). Synaptic puncta, identified by staining for vGlut1 and synapsin (SYN1), were observed on TUJ1- or MAP2-neurites at day 35 (Figure 1H, 1I, and S1I). Collectively, the fibrin gel supports the maturation and synaptogenesis of the cortical neurons.

### Printed neural cells mature and retain the tissue structure

A basic requirement for printing functional neural tissues is to enable neuronal maturation while maintaining the tissue structure. We printed the cell layer, or “band” of ∼50 µm thickness horizontally, one after another. These horizontal “bands”, when turned 90°, exhibit vertical “layers” (Figure 2A). To better visualize our printed tissue construct, the GFP- or mCherry-labeled NPCs were loaded in the bioink and printed one band at a time at the dimension of 5000 (L) × 500 (W) × 50 (H) µm (Figure 2A; Video S1). To prevent the mix of the printed bands, we added the crosslinking agent-thrombin immediately following the deposit of the cell-gel mixture to form the desired shape before printing the next band (Figure 2A; shape design based on the Supplementary G.code files). The printed neural tissue constructs were generated with multi-layered patterns (Figure 2B and S1J). Indeed, the GFP^+^ cells in one band were well separated from the mCherry^+^ cells in the next while becoming MAP2^+^ maturing neurons at 7 days after printing (Figure 2C). Although the printed cells stayed within the designated areas, the neurons extended processes in and across the bands (Figure 2D and S1K). There were multiple cells in depth of the 50 µm-thick tissues (Figure 2D; Video S2), which was maintained by the conventional culture system and monitored by conventional microscopy. Thus, the printed tissue retains a stable structure within which the neural progenitors mature and form neural networks.

**Figure 2.**
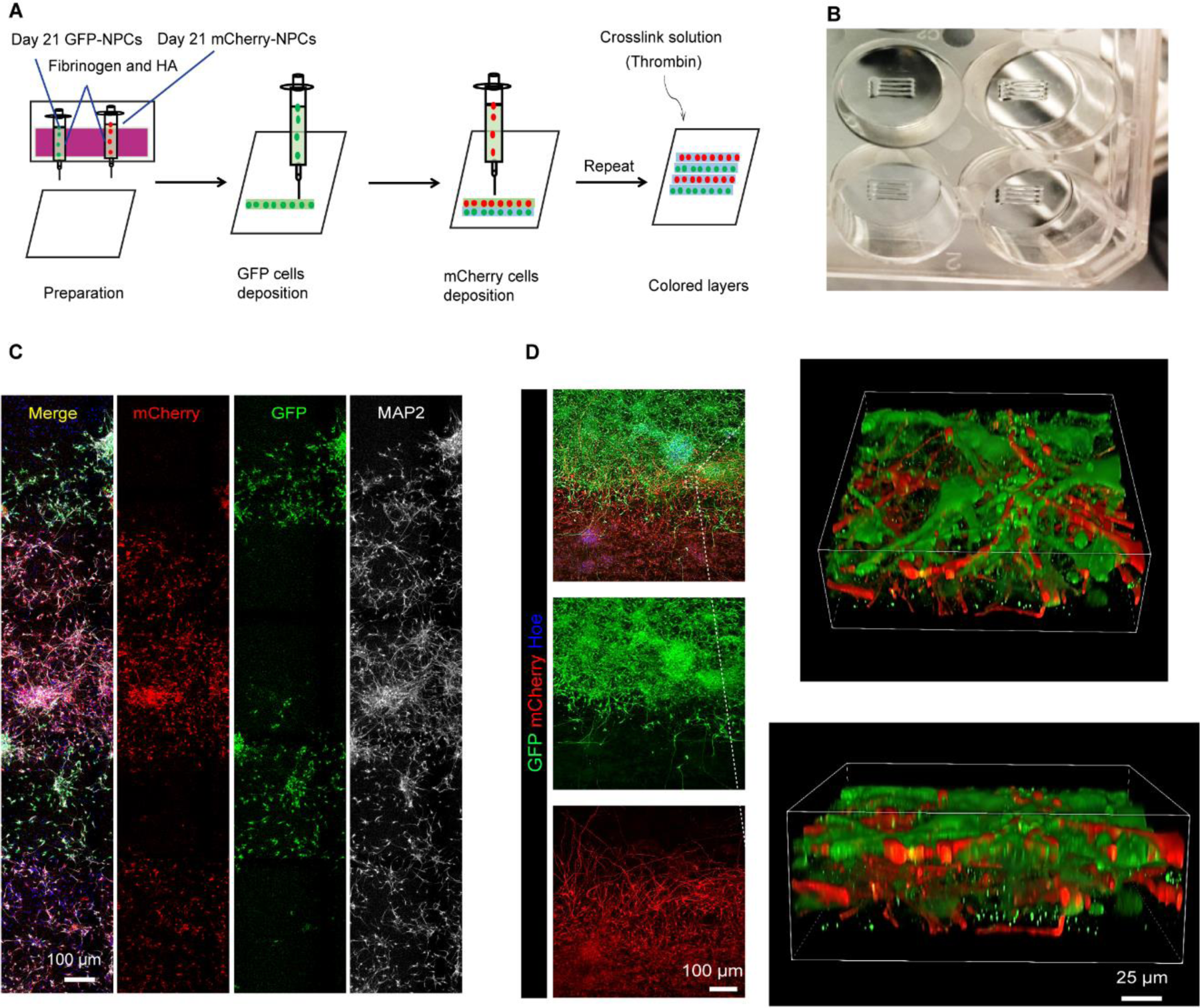
Printing neural tissues by design. (A) Design of the printing process for layer-patterned neural tissues. See Video S1 shows the printing process. HA, hyaluronic acid. (B) Overview of the printed structure (5000 × 500 × 50 µm) using the testing bioink in a 24-well plate. (C) Printed neural tissue with the “green-red-green-red” layered pattern 7 days post-printing immunostained for MAP2. (D) Printed neural tissue with a “green-red” layered pattern (15 days post-printing), immunostained for GFP and mCherry. The inset is enlarged with a 3D reconstruction view. See Video S2 shows 3D reconstruction of printed neurons. Blue color, Hoechst 33342 staining. Hoe, Hoechst 33342. See also Figure S1 and Table S2.

### Neuronal subtypes form functional networks in printed tissues

The brain functions through interactions between different neuronal types. In the cerebral cortex, two major neuronal types, GABAergic interneurons and glutamatergic neurons, synapse and interact with each other. To determine if these two neurons, incorporated into the printed tissues, form synaptic connections, we generated medial ganglionic eminence (MGE) GABA and cortical (glutamate) progenitors from GFP^+^ and GFP^-^ hPSCs and mixed the two progenitor populations at a ratio of 1:4 before printing to mimic the ratio of interneurons and cortical projection neurons in the cerebral cortex (Tremblay et al., 2016; Wonders and Anderson, 2006) (Figure 3A). The hPSC-derived MGE cells at day-21 expressed NKX2.1, GABA and negative for PAX6 (Figure S2A-S2C), whereas the cortical progenitors were positive for PAX6 and FOXG1 (Figure S1A). To show the maintenance of the layered structure while examining the neuronal connections within and between the bands, we printed a two-layered tissue, in which one band was GFP^+^ cells and the other was non-colored cells, and each band contained the same proportion of interneurons and cortical projection neurons (Figure 3A). The printed tissue with only cortical projection neurons was served as a reference. As shown in Figure 3B, both the GFP^+^ and GFP^-^ bands displayed an even distribution of GABA^+^ neurons and all MAP2^+^ neurons. The GABA^+^ cells comprised about 20% of total MAP2^+^ neuronal population at day 3 post-printing and this ratio was maintained at day 20 (Figure 3C, 3D, S2D, and S2F). As a reference, few GABA^+^ interneurons (less than 1.5%) were found in the tissue printed with cortical progenitors only (Figure S2E and S2F). Hence, the neuronal subtypes in the printed tissue are maintained over time.

**Figure 3.**
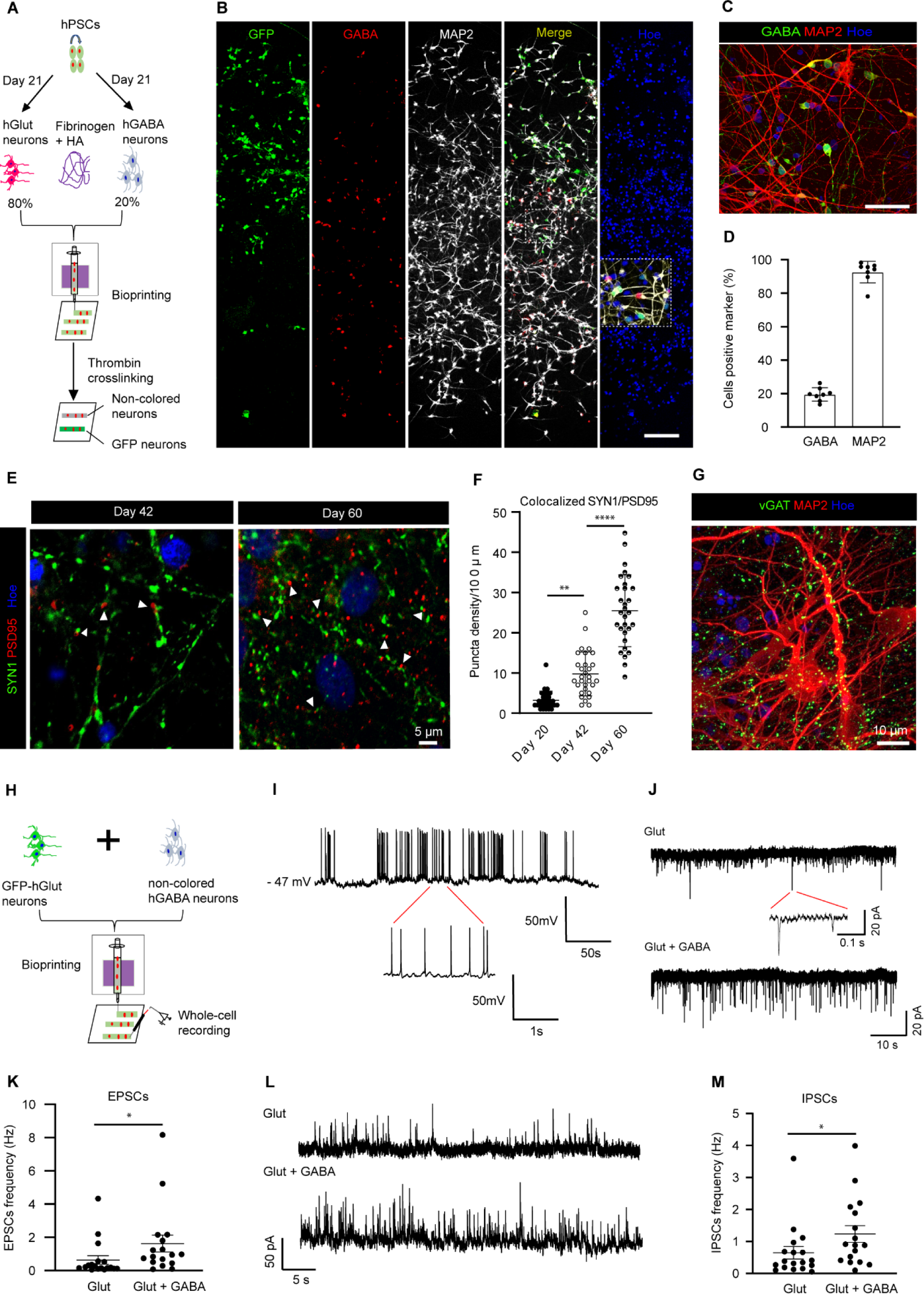
Functional connections between neurons in the printed tissue. (A) Schematic diagram illustrating the design of experiments. (B) Immunostaining of the tissue for GABA and MAP2 (3 days post-printing) (Scale bars, 100 µm). (C) Immunostaining of tissues for GABA and MAP2 (20 days post-printing) (Scale bars, 50 µm). (D) Quantitative analysis of GABA and MAP2 cell populations (8 samples from 3 three different batches were analyzed). (E) Immunostaining for synaptic markers SYN1 and PSD95 at day 42 and 60. White triagnles indicate the co-localized punctas of SYN1 and PSD95. (F) Quantitative analysis of colocalized SYN1 and PSD95 puncta density over time (one-way ANOVA, **P < 0.01, ****P < 0.0001, 30 neurons from three different experiments were analyzed). (G) Immunostaining for synaptic markers vGAT with MAP2 at day 30. (H) Schematic diagram illustrating the electrophysiological recording of printed tissues. (I) Spontaneous action potentials (sAPs) of glutamate cortical cells from printed tissues incorporated with Glut and GABA neurons at day 21 post-printing. (J) Representative traces of EPSCs recorded from glutamatergic neurons in tissues with Glut neuron-only or Glut and GABA (Glut + GABA) tissues at 5 weeks post-printing. (K) Quantitative analysis of EPSC frequency for different groups 5 weeks after printing (Mann-Whitney U-test, *P < 0.05; 18 neurons from five different batches were analyzed for Glut condition and 17 neurons from five different batches were analyzed for Glut + GABA condition). (L) Representative traces of IPSCs recorded from glutamatergic neurons in printed tissues with Glut neuron-only or Glut and GABA neurons (Glut + GABA) at 5 weeks post-printing. (M) Quantitative analysis of IPSC frequency for different groups 5 weeks after printing (Mann-Whitney U-test, *P < 0.05; 18 neurons from five different batches were analyzed for the Glut group and 17 neurons from five different batches were analyzed for the Glut + GABA group). Data are represented as mean ± SEM. Hoe, Hoechst 33342. See also Figure S2-S4.

Examination of tissues printed from the non-colored hPSCs (Figure S3A) at a longer time post-printing (day 60) indicated that the cortical neurons expressed cortical transcription factors TBR1, CTIP2 and SATB2 (Figure S3B). We also observed GABAergic subtypes, labeled by calretinin (CR), calbindin (CB), parvalbumin (PV) and somatostatin (SST) (Figure S3C). Therefore, the printed neural tissue constructs support the maturation of cortical glutamatergic neurons and GABAergic interneurons.

Neuronal maturation was reported in printed 3D human neural tissues showing expression of MAP2 and NeuN (Table S1). We found the co-expression of the pre- and post-synaptic markers, SYN1 and PSD95, as early as day 20 after printing (Figure 3E), indicating the formation of synapses, which usually needs longer time to form in 2D cultures (Gunhanlar et al., 2017; Shi et al., 2012). The synaptic puncta density increased over extended culture (Figure 3E and 3F). More SYN1 puncta density was present in the printed tissues with both glutamate and GABA neurons than in prints with only glutamatergic neurons (Figure S3D and S3E), suggesting that interactions between glutamatergic neurons and GABAergic neurons promote synaptogenesis. The GABAergic neurons also formed synapses as indicated by the expression of inhibitory presynaptic protein vGAT (Figure 3G). Furthermore, the excitatory pre-synaptic marker vGlut1 and inhibitory postsynaptic marker Gephyrin were expressed at day 25 after printing, and more vGlut1 and Gephyrin puncta were observed at day 45 (Figure S3F). These observations indicate the formation of excitatory and inhibitory synapses in the printed neural tissues.

The most challenging aspect of human neural tissue printing is formation of functional networks. To date, few 3D-printed human neural tissues show functional connectivity except for cells seeded in the pre-set scaffolds or molds (Table S1). Given the appropriate thickness of printed tissue construct, we performed electrophysiological recording of the tissues that were printed with GFP^+^ glutamatergic cortical progenitors and non-colored MGE GABAergic progenitors (Figure 3H). Whole-cell patch clamping indicated that the glutamatergic neurons exhibited inward Na^+^ and outward K^+^ currents (Figure S4A-S4C) and evoked action potentials (eAP) about 3 weeks after printing (Figure S4D). And these neurons also exhibited spontaneous action potentials (sAP) (Figure 3I). The percentage of cortical neurons with sAP increased overtime, particularly in the prints with the incorporation of GABAergic neurons (36.36% vs 47.06 % for 3 weeks and 50% vs 71.43% for 5 weeks) (Figure S4E). The frequency of sAP also increased with time and printed tissues with GABAergic and glutamatergic neurons showed more sAP (Figure S4F and S4G), suggesting functional maturation of the printed neurons. Importantly, the glutamatergic neurons displayed excitatory and inhibitory postsynaptic currents (EPSCs & IPSCs), suggesting formation of functional networks between glutamatergic neurons and GABAergic interneurons (Figure 3J and 3L). Moreover, these cells received more EPSCs and IPSCs when printing together with GABAergic neurons (Figure 3K and 3M). The hPSC-derived cortical GABAergic interneurons usually mature very slowly (Nicholas et al., 2013). In the printed tissue, the GABAergic interneurons generated spontaneous action potentials at day 21 (Figure S4H). Spontaneous IPSCs were observed in the GABA neurons at day 35 and the IPSCs were abolished by the GABA_A_ receptor antagonist bicuculline (Figure S4I), indicating that GABA neurons were also functionally mature. These results confirm the functional maturation of cortical projection neurons and GABAergic interneurons and the functional connection among neurons within the printed tissues.

### Neurons and astrocytes form functional networks in printed tissues

Appropriate neuronal network function requires the presence of glia, including astrocytes. We hence incorporated hPSC-derived astrocyte progenitors (Figure S5A and S5B) into the above glutamate neurons and GABA interneurons at a ratio of 4:5:1 within the printed tissue. We also printed a two-layered tissue consisting of GFP^+^ and GFP^-^ cells; and each band had the three cell types (Figure 4A). The GFAP^+^ astrocytes and the MAP2^+^ neurons were distributed throughout the printed tissue (Figure 4B), as were GABA^+^ interneurons (Figure S5C). Quantitative analysis of GABA, MAP2 and GFAP expression showed that the proportion of the three cell types in printed neural tissues was maintained (Figure S5E).

**Figure 4.**
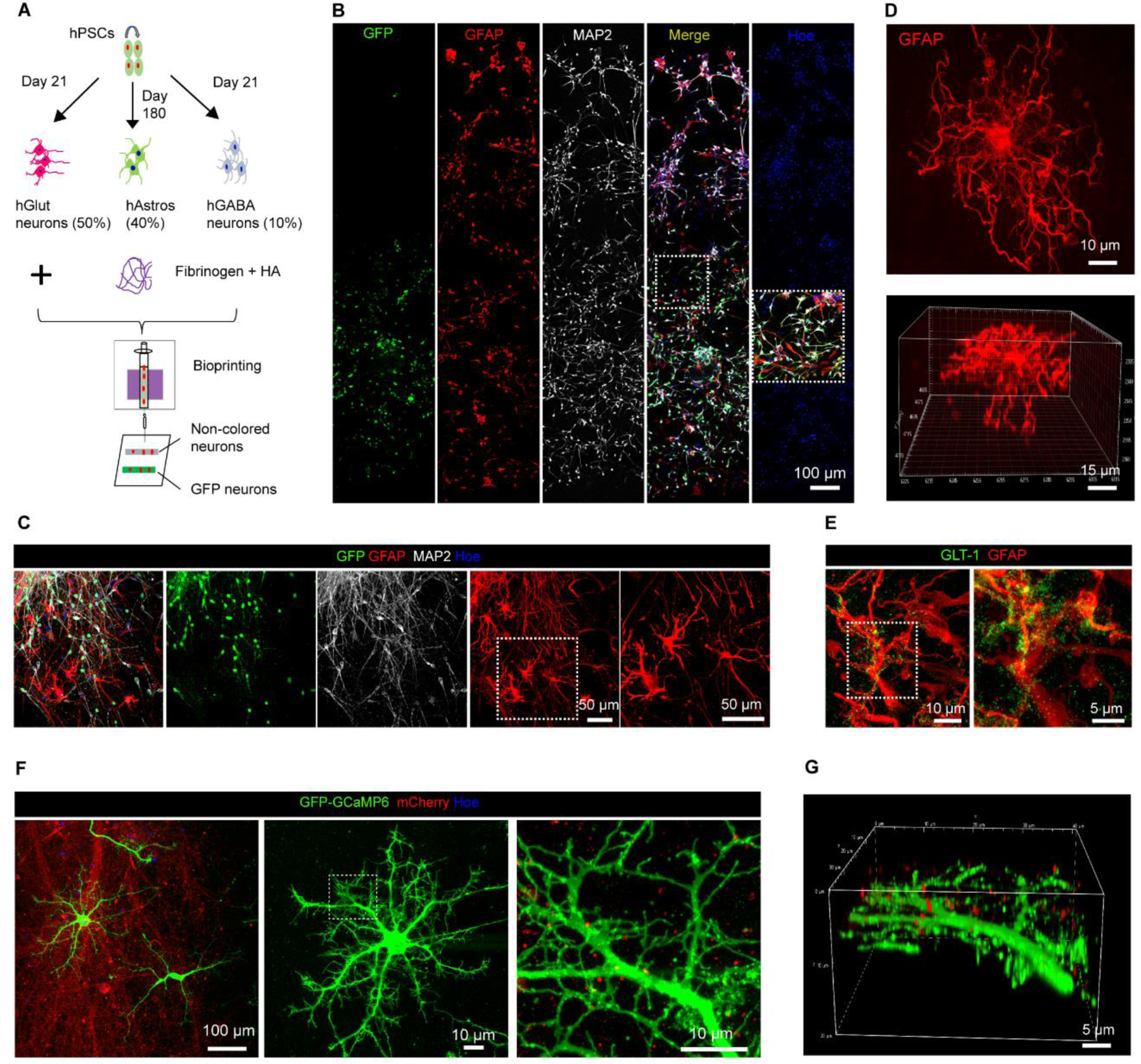
Incorporation of astrocytes with neurons in printed tissues. (A) Schematic diagram illustrating the design of experiments. (B) Immunostaining of the tissue for GFAP and MAP2 (3 days post-printing). Hoe, Hoechst 33342. (C) Immunostaining of astrocytes and neurons in printed tissues for GFAP and MAP2 at day 60 post-printing. Hoe, Hoechst 33342. (D) The 3D morphology of astrocytes with staining of GFAP in printed tissues at day 60. (E) Immunostaining of tissues for GLT-1 and GFAP at day 40. (F) Immunostaining for GFP-GCaMP6 and mCherry at day 30 post-printing. (G) 3D imaging from (F). See also Figure S5 and S6.

In the prints with both neurons and astrocytes, most of the neurons (> 90%) became NeuN^+^ neurons at day 30 post-printing (Figure S6D and S6E). Particularly, we discovered that astrocytes became progressively more mature over time with more elaborate processes and branches (Figure 4C, 4D, S6B, and S6C). While GFAP staining showed the extensive branches of astrocyte processes (Figure 4D), GFP labeling revealed fine processes and their sheets (Figure 4F, S6I, and S6J). Notably, 3D visualization showed that the astrocytic branches were intertwined with the mCherry-neuronal processes (Figure 4G and S6J), suggesting close neuron-astrocyte interactions.

Astrocytes, in response to neuronal stimulation, generate calcium signals, which enables them to modulate neuronal networks (Ben Haim and Rowitch, 2017; Khakh and Sofroniew, 2015). Neurons and astrocytes were printed together into neural tissues (Koroleva et al., 2021; Salaris et al., 2019; Zhou et al., 2020) (Table S1), but whether they form functional connections is not known. To determine if the astrocytes functionally integrate into the printed neural networks, we printed mCherry-neurons with lentivirus-GCaMP6 infected astrocytes (Figure 5A). We found that the application of a high concentration KCl solution, which depolarizes neurons but does not obviously affect astrocytes directly, elicited calcium responses in astrocytes in the printed tissues (Figure 5B and 5C). The total *ΔF/F* of GCaMP6 signals increased from 2 to 6 weeks after printing (Figure 5D). Thus, the neurons and astrocytes in the printed tissues are functionally connected.

**Figure 5.**
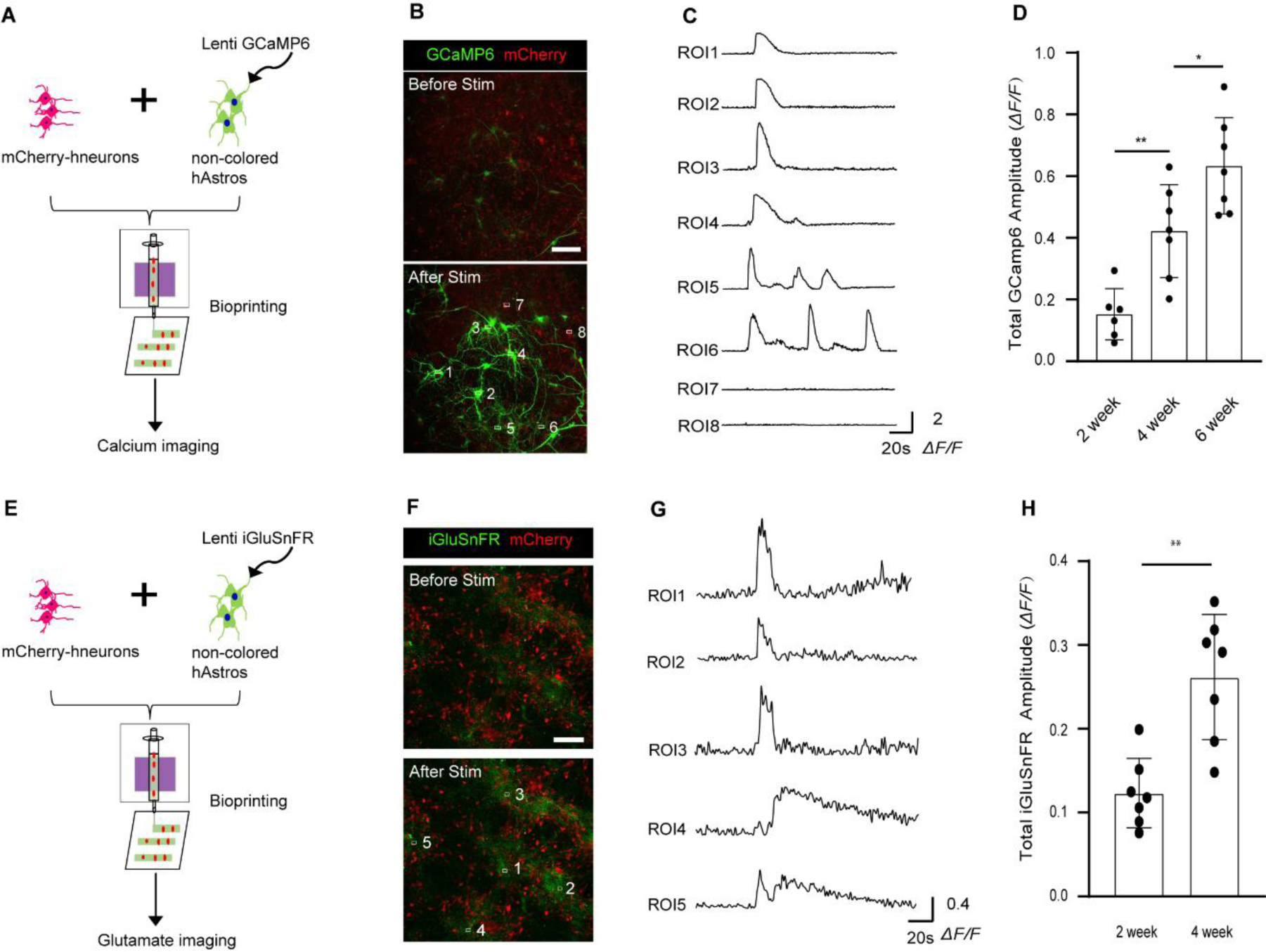
Neural network between neurons and astrocytes in the printed tissue. (A) Schematic diagram illustrating calcium imaging of printed tissues with neurons and astrocytes. Neurons were mCherry labeled and astrocytes were infected with lentivirus-GCaMP6 before printing. (B) Cells in printed tissues responding to neuronal stimulation with a high concentration of KCl solution (Scale bars, 100 µm). (C) Representative traces of GCaMP6 imaging. ROI1-6 indicate GCaMP6-astrocytes and ROI7, 8 indicate mCherry-neurons. ROI, region of interest. (D) Total GCaMP6 amplitudes of *ΔF/F* in tissues at 2, 4, and 6 weeks after printing (one-way ANOVA, **P < 0.01; *P < 0.05, 6 -7 samples from three different batches were analyzed for each time point). (E) Schematic diagram illustrating glutamate imaging of printed tissues with neurons and astrocytes. Neurons are mCherry labeled and astrocytes were infected with lentivirus-iGluSnFR before printing. (F) Cells in printed tissues responding to neuronal stimulation with high concentration KCl solution (Scale bars, 100 µm). (G) Representative traces of iGluSnFR imaging. ROI1-5 indicate iGluSnFR-astrocytes. (H) Total iGluSnFR amplitudes of *ΔF/F* in tissues at 2 and 4 weeks after printing (t-test, **P < 0.01, 7 samples from three different batches for each time point). Data are represented as mean ± SEM.

One of the most important functions of astrocytes is to recycle neurotransmitters like glutamate that are released to the synaptic cleft (Ben Haim and Rowitch, 2017). Indeed, the astrocytes expressed GLT-1 (glutamate transporter 1) at day 40 post-printing (Figure 4E), suggesting maturation of astrocytes. We further examined networks between neurons and astrocytes using live imaging of a glutamate indicator iGluSnFR (Figure 5E). Again, when applying KCl solution, which triggers cortical neurons to release glutamate but does not have an obvious effect on astrocyte-only cultures, the iGluSnFR signals in astrocytes were induced (Figure 5F and 5G). The total *ΔF/F* of iGluSnFR signals increased from 2 to 4 weeks after printing (Figure 5H). Therefore, astrocytes can take up glutamate released from neurons in the printed tissues.

### Functional networks form between the printed cortical-striatal tissue layers

Neurons not only synapse each other in the same brain region but also connect to each other between nuclei or layers. Neural tissues printed in a thick stack do not project neurites (nerves) across the layers (Table S1). To assess the functional connectivity between tissue layers, we printed cortical neurons with DARPP32^+^ striatal medium spiny neurons (Figure 6A). The striatal neurons showed high expression of GABA and DARPP32 (Figure S7A-S7D). After printing, cortical neurons expressed GFP, and striatal neurons, expressing mCherry and DARPP32, showed a distinguished tissue separation (Figure S7E). Both neuronal types became mature with expression of MAP2 at 2 weeks after printing (Figure S7F). The printed cortical and striatal neuronal bands were well-maintained 15 days after printing, and GFP and mCherry neurites grew toward each other though the GFP-labeled cortical neurites projected deeper into the mCherry-labeled striatal band (Figure 6B). The projected cortical neurites (axons) formed physical contacts with the striatal neurites (Figure S7G; Video S3), indicating the formation of connection.

**Figure 6.**
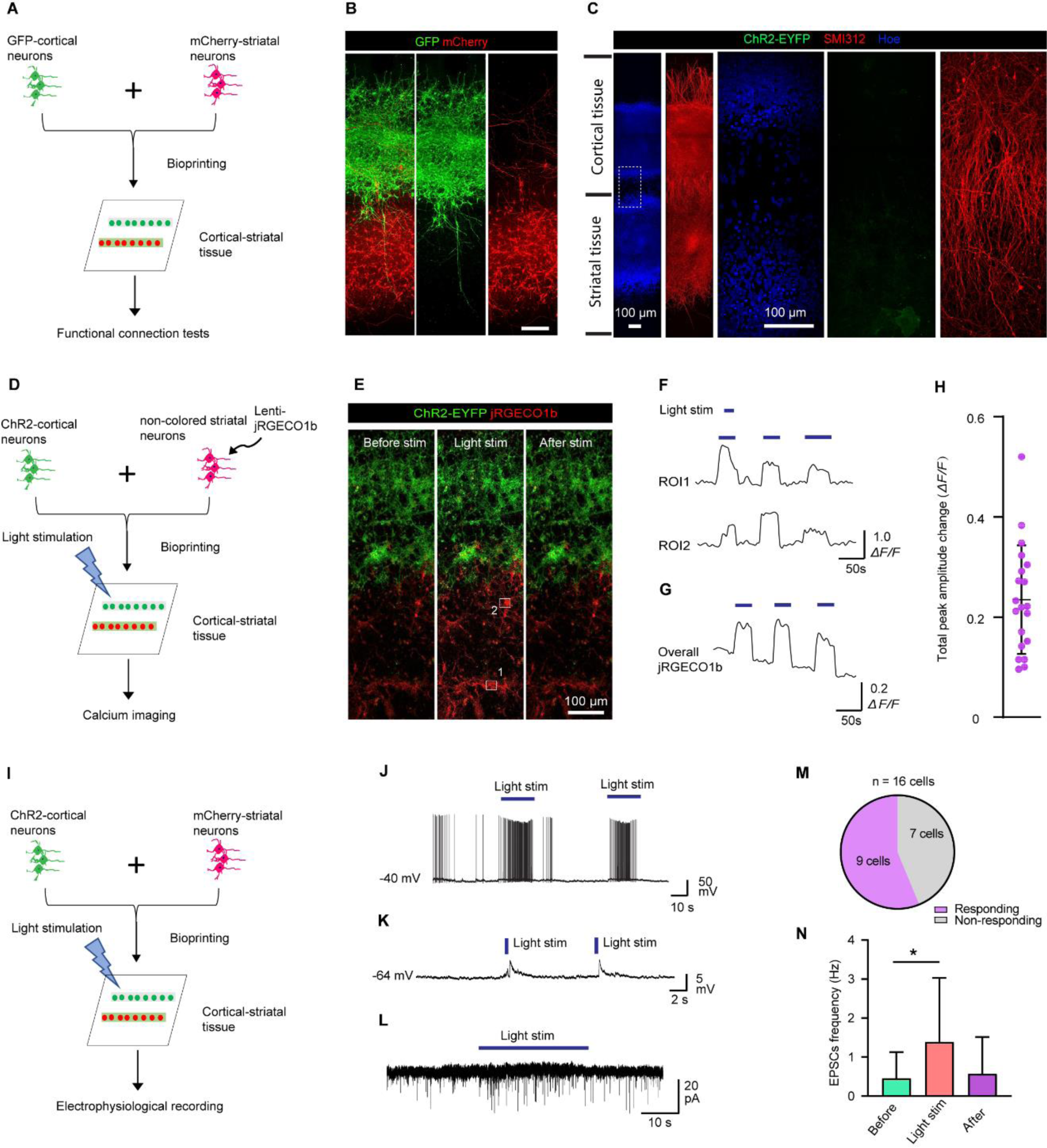
Functional network between cortical and striatal neuronal layers. (A) Schematic diagram illustrating the design of experiments for printing the cortical-striatal tissue. (B) Immunostaining of the tissue for GFP and mCherry to show the printed cortical-striatal layers at 15 days post-printing (Scale bar, 100 µm). (C) Immunostaining of cells for the axonal marker SMI312 in the 3D printed tissue (4 week after printing). (D) Schematic diagram illustrating calcium imaging of printed tissues with cortical and striatal neurons. Cortical neurons expressed ChR2, and striatal neurons were non-colored and infected with lentivirus-jRGECO1b before printing. (E) Printed tissue (2 weeks post-printing) responding to light stimulation, showing changes in jRGECO1b signals (red) before and after stimulation. (F) Representative traces of calcium imaging from two regions of interest. ROI, region of interest. (G) Total traces of calcium imaging. (H) Total jRGECO1b amplitudes of *ΔF/F* in tissues after light-stimulation (21 samples from five different batches). (I) Schematic diagram illustrating the method for whole-cell patch-clamp recording with optogenetic activation in printed cortical-striatal tissues. Cortical neurons were ChR2 cells, and striatal neurons were mCherry labeled. (J), (K) and (L) Representative electrophysiological traces of spontaneous action potentials, EPSPs and EPSCs of striatal neurons within printed tissues by light stimulation. (M) Percentage of responsive cells with light stimulation for EPSCs recording. (N) Quantitative analysis of EPSC frequency before, during and after light stimulation (paired t-test, *P < 0.05; 9 neurons from three different batches). Data are represented as mean ± SEM. Hoe, Hoechst 33342. See also Figure S7.

In the brain, cortical neurons (deep layers) project axons to the striatum. In our print, we observed neurites growing toward each other (Figure 6B). To determine if the cortical and striatal neuron axons project to each other, we performed the immunostaining of the printed tissue with an axonal marker SMI312 when the cortical-striatal tissue was printed with a gap of 100 µm between the bands (Figure 6C). Strikingly, we found that the axonal projection was from the cortical neurons to striatal neurons but not from the striatal to cortical neurons (Figure 6C). This pattern of axonal projection resembles that in vivo where cortical neurons project axons to the striatum, demonstrating the striking specificity of neuronal network formation in the printed tissues.

To determine whether neurons in the printed cortical tissue form functional synaptic connections with those in the striatal tissue, we printed cortical neurons that expressed ChR2-EYFP (Dong et al., 2020) with striatal neurons that expressed the red calcium indicator jRGECO1b (Dana et al., 2016) (Figure 6D). We found that application of 470-nm light to stimulate ChR2-EYFP cortical neurons elicited calcium responses in the striatal neuron band 2 weeks after printing (Figure 6E). Even the striatal neurons far from the cortical band showed increased calcium responses (Figure 6F). The total ΔF/F of calcium signals for the whole striatal neuron layer significantly increased with stimulation (Figure 6G and 6H), suggesting functional connections of the cortical neuron band with the striatal neuron band. We further characterized the functional connections between cortical and striatal neurons by patch clamping in response to light stimulation (Figure 6I, S7H, and S7I). We found that the application of 470-nm light induced burst action potentials, spontaneous excitatory postsynaptic potentials (EPSPs) in the striatal neurons 3-5 weeks after printing (Figure 6J-6L). The percentage of responsive cells for EPSCs was > 56% (Figure 6M). And the frequency of EPSCs was significantly increased by light stimulation (Figure 6N and S7I). Thus, both the calcium imaging and electrophysiological recording demonstrate the functional connectivity between the cortical-striatal tissue layers.

As shown above, stimulation of the ChR2-expressing cortical neurons elicited response in the striatal neurons as assayed by calcium signaling (Figure 6D-6H). The question is whether stimulation of striatal neurons elicits response in the cortical neurons. We printed ChR2-expressing striatal neurons and jRGECO1b infected cortical neurons (Figure S7J). At 2 and 4 weeks after printing, light stimulation of the ChR2-expressing striatal neurons did not elicit obvious response in the cortical neurons based calcium imaging (Figure S7K and S7L). The lack of response from the cortical neurons to stimulation of striatal neurons is likely due to the lack of axonal projection from the striatal neurons to cortical neurons. This finding again demonstrates the specificity of functional connections between the cortical-striatal tissues in the prints.

### The printed human neural tissues are amenable for modeling pathological processes

To assess if the above printed tissues are amenable for examining pathological processes, especially at the functional level, we use Alexander disease (AxD), a neurodegenerative disease caused by mutations in the glial fibrillary acidic protein (GFAP) gene (Messing et al., 2012), as an example. We assessed neuron-astrocyte interaction in the prints similar to that described in Figure 4A. In this case, the (glutamate and GABA) neurons were derived from hESCs (H9) whereas the astrocytes were derived from either AxD (R88C mutation) or isogenic control (mutation corrected by CRISPR) iPSCs (Jones et al., 2018) (Figure 7A). After printing, the AxD astrocytes displayed GFAP aggregation intracellularly compared to the isogenic controls (Figure 7B), which recapitulates the pathological feature of AxD (Mignot et al., 2004). The printed AxD tissues showed less expression of GLT-1 comparing to the control at day 20 post-printing (Figure 7C). By 30 days after printing, MAP2^+^ neurons and GFAP^+^ astrocytes displayed a complex morphology with elaborate processes and expression of synapsin (Figure 7D). Interestingly, the AxD tissues had a significantly reduced synaptic puncta density than the tissue with isogenic astrocytes (Figure 7D and 7E), suggesting that AxD astrocytes are less supportive for synaptogenesis compared to the control astrocytes.

**Figure 7.**
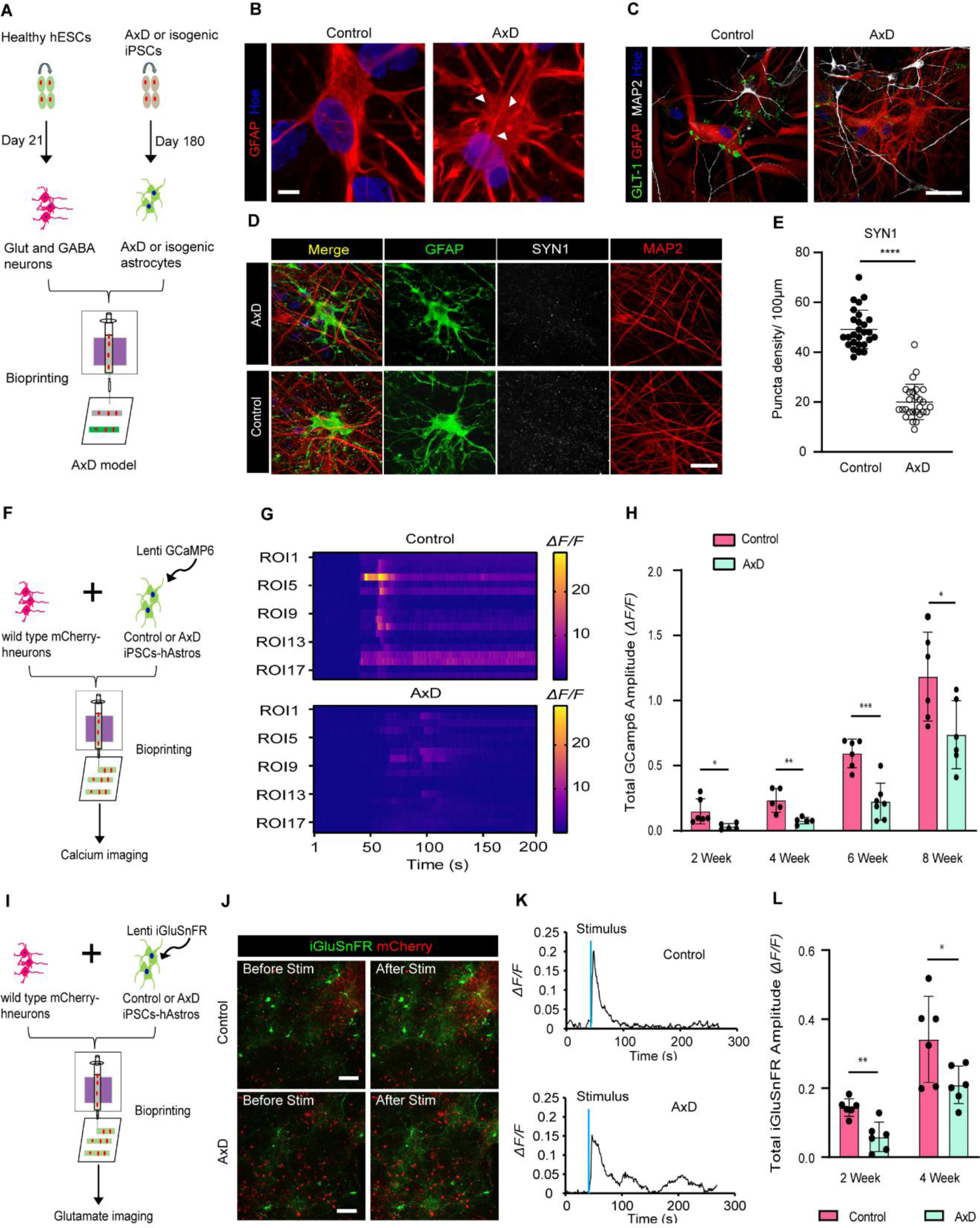
Modeling AxD using the printed tissue. (A) Schematic diagram illustrating the design of experiments. We used a pair of AxD patient-iPSCs and its isogenic control. (B) Immunostaining of control and AxD tissues for GFAP showing the aggregation of GFAP fibers at day 60 post-printing. White triangles indicate the GFAP aggregates. Similar results were observed in three independent experiments. (C) Immunostaining of control and AxD tissues for GLT-1, GFAP and MAP2 at day 20 post-printing. (D) Immunostaining of tissues for SYN1, GFAP and MAP2 (30 days post-printing). (E) Quantitative analysis of SYN1 puncta density (t-test, ****P < 0.0001, 30 samples for each condition from at three different experiments). (F) Schematic diagram illustrating calcium imaging of printed tissues for AxD model. Healthy neurons were mCherry-labeled and AxD or control astrocytes were infected with GCaMP6 before printing. (G) Heatmap showing *ΔF/F* of GCaMP6 signal from control and AxD. (H) Total GCaMP6 amplitudes of *ΔF/F* in control and AxD tissues overtime (multiple t-tests, *P < 0.05; **P < 0.01; ***P < 0.001, n = 6 from three different experiments). (I) Schematic diagram illustrating glutamate imaging of printed tissues for the AxD model. Healthy neurons were mCherry-labeled and AxD or control astrocytes were infected with iGluSnFR before printing. (J) Cells in printed tissues responding to neuronal stimulation with a high concentration KCl solution. (K) Representative traces of iGluSnFR change (*ΔF/F*) from the control and AxD tissues. (L) Total iGluSnFR amplitudes of *ΔF/F* in the control and AxD tissues at 2 and 4 weeks after printing (multiple t-tests, *P < 0.05; **P < 0.01, n = 6 from three different experiments). Data are represented as mean ± SEM. Hoe, Hoechst 33342. Scale bars, 5 µm (B), 20 µm (D), 50 µm (C), 100 µm (J).

We have previously shown that AxD patient astrocytes in 2D cultures failed to propagate calcium waves (Jones et al., 2018). To determine if AxD astrocytes alter their response to neuronal function, we used the prints of normal neurons with either AxD or isogenic astrocytes infected with lentivirus-GCaMP6 (Figure 7F). Stimulating neurons with high KCl elicited a robust Ca^+^ response in isogenic astrocytes. In contrast, AxD astrocytes showed significantly fewer calcium responses (Figure 7G). The total *ΔF/F* of GCaMP6 signals for AxD astrocytes was significantly lower over time compared to that of control (Figure 7H). We further investigated the neuron-glial connections in AxD using live imaging of glutamate uptake by iGluSnFR (Figure 7I). Upon neuronal stimulation with high KCl, the AxD astrocytes in printed tissues exhibited fewer changes of iGluSnFR signals comparing to the isogenic control (Figure 7J and 7K). The total *ΔF/F* of iGluSnFR signals for AxD astrocytes was significantly lower from 2 and 4 weeks after printing as compared to that of the isogenic control (Figure 7L). These results indicate that the disease relevant functional phenotypes are readily displayed in the printed human neural tissues.

## DISCUSSION

We have printed functional human neural tissues by design. We achieve it by first developing a bioink that is suitable for the survival, neurite growth and synapse formation by the differentiating neurons. Then we designed a special tissue pattern by printing the bands horizontally, allowing easy live-cell imaging and electrophysiological recording. Importantly, the printed neural tissues form functional synaptic connections between neuronal subtypes in and between tissue bands as well as functional neuron-astrocyte networks within 2-5 weeks after printing their progenitors. The brain tissues printed by design offer a defined platform for examining human neural networks under physiological and pathological conditions.

Our platform has several features. First, the functional neural circuits, indicated by EPSCs and IPSCs, are established in 2-5 weeks after printing. Multi-layered neural tissues have been fabricated through vertical deposition (Edelbrock et al., 2021; Joung et al., 2018; Kajtez et al., 2022; Lozano et al., 2015; Skylar-Scott et al., 2022), which requires stiff bioink to hold the tissue stable. These printed tissues show little functional connectivity possibly due to the stiff bioink, as only soft gels support nerite sprouting and synapse formation (Georges et al., 2006; Koser et al., 2016). Second, the printed tissue allows the incorporation of different neuronal subtypes such as cortical glutamatergic neurons, cortical GABAergic interneurons or striatal medium spiny neurons at defined proportions, which is difficult to achieve via organoids or other printing methods to date. Third, this platform enables establishment of functional neuron-glial networks with a controlled neuron-to-glia ratio within 2-4 weeks, simplifying the analysis of neuron-glial interactions in the 3D environment. Fourth, functional layered tissues like the cortical-striatal tissue can be readily assembled and the intrinsic property of tissue interaction such as axonal projection is retained, as indicated by the cortical projection to the striatal tissue. To our knowledge, this is the first to assemble region-specific brain tissues with functional connectivity between them using a bioprinting system. Interaction between different brain regions may also be achieved using assembloids though the assembloids are often combined in a random fashion and it takes much longer time for them to form functional connections (Andersen et al., 2020; Miura et al., 2020; Xiang et al., 2017). Our tissue assembly can be printed in a defined orientation and distance. Such a printing platform provides a promising model to study human neural circuits among different brain regions. Fifth, the dimensions of the printed tissues, including the thickness, are controlled so that they allow sufficient nutrients and oxygen for the survival, growth, and function of the neural tissue under the conventional culture system. This relies on our special design of horizontal printing, overcoming thickness of the vertical printing and the spheric structure of organoids. Finally, the composition of the functional cell types depends on that of printed progenitor cells, making it predictable for the anatomical and potentially functional properties of the printed tissues. Together, these features make the printed neural tissue a promising paradigm for studying human neural network functions.

### Limitations of the study

Our prototype 3D bioprinting has weaknesses. Due to the softness of the gel, our bioink does not support multiple layer printing vertically. We also limited the thickness of the printed tissues to about 50μm to maximize the formation of functional neural networks. Additionally, our current printing technology does not enable the orientiation of the mature neurons even though many neurons exhibit a pyramidal morphology. Since the tissue is assembled by design, the printed tissue lacks the intrinsic structural organization as brain oragnoids. Interestingly, however, some of the intrinsic properties, including the anatomical and functional neuronal connections, are retained in the printed tissue, as evidenced by the uni-directional axonal projection from the cortical tissue to the striatal tissue. On the other hand, the cells in the printed brain tissue mature rapidly and in a synchronized mammner, thus complementing the existing organoids and offering a defined platform for examining human neural networks under physiological and pathological conditions. It is expected that with advancement of the bioprinting technology, more sophisticated human neural tissues may be produced with defined cellular compositions and orientation, tissue organization, and tissue assembly. Given that many neuronal subtypes can now be generated from hPSCs (Tao and Zhang, 2016), our platform allows printing neural tissues with a defined composition of neural types at a particular ratio, potentially enabling assessment of the biophysical nature of the human neural circuits. As exemplified by the dynamic interaction between neuronal activation and altered response from AxD astrocytes in the printed neural tissues, the platform also provides a promising tool for studying interactions between specific neural cell types and neural circuits under pathological conditions. The defined dimensions and cellular compositions make the platform amenable for throughput analysis and for drug development.

## Supporting information

Resource table

## ACKNOWLEDGEMENTS

We thank X. Chu, and M. Ayala for technical assistance; K. Knobel for confocal imaging and Imaris image analysis. This study was supported in part by NIH-NINDS (NS096282, NS076352, NS086604), NICHD (HD106197, HD090256), the National Medical Research Council of Singapore (MOH-000212, MOH-000207), Ministry of Education of Singapore (MOE2018-T2-2-103), the Bleser Family Foundation, and the Busta Foundation. This research was also funded in part by Aligning Science Across Parkinson’s (ASAP-0031) through the Michael J. Fox Foundation for Parkinson’s Research (MJFF). For the purpose of open access, the author has applied a CC BY public copyright license to all Author Accepted Manuscripts arising from this submission.

## AUTHOR CONTRIBUTIONS

Y.Y. conceived and designed the study, performed the cell culture, differentiation, immunostaining, bioink preparation, printing, calcium and glutamate imaging, data analysis and interpretation, and wrote the manuscript. X.L., S.M. & Y.D. performed the electrophysiology. L.K. provided astrocytes and produced GCaMP6 virus. Y.G. & X.Z. produced iGluSnFR virus. Y.T., X.L. & A.B. performed data interpretation. SC.Z. conceived and designed the study, acquired the funding, supervised the study, performed data analysis and interpretation, and wrote the manuscript.

## STAR ★ METHODS

### RESOURCE AVAILABILITY

#### Lead contact

Further information and requests for resources and reagents should be directed to and will be fulfilled by the lead contact, Dr. Su-Chun Zhang.

#### Materials availability

This study did not generate new unique reagents.

#### Data and code availability

The G.code for the design of print construct is available in the Supplemental files.

### EXPERIMENTAL MODEL AND SUBJECT DETAILS

#### Human cell source

Human ESCs (H9, GFP-H9, mCherry-H9, and ChR2-EYFP) and iPSCs (AxD R88C and isogenic control) were maintained on the mouse embryonic fibroblast (MEF) feeder in a stem cell growth medium or on Matrigel-coated plates in the TeSR-E8 medium (StemCell Technologies, Inc., Vancouver, Canada) as described previously (Li et al., 2018; Yan et al., 2015). For MEF feeder-based cultures, cells were passaged weekly by using dispase (1 mg/mL) and plating on a monolayer of irradiated MEF (WiCell). The hPSC culture medium consisted of DMEM/F12 basal medium (Gibco), 20% KnockOut serum replacement (Gibco), 0.1 mM β-mercaptoethanol (Sigma), 1 mM L-glutamine (Gibco), nonessential amino acids (Gibco), and 4 ng/mL fibroblast growth factor (FGF)-2 (R&D Systems). For TeSR-E8 medium-based cultures, cells were passaged every 6-7 days by accutase and plated on Matrigel-coated 6-well plates for monolayer cultures in the presence of ROCK inhibitor Y27632 (10 µM) to promote cell survival.

### METHOD DETAILS

#### Generation of cortical neural progenitors, GABAergic interneurons, striatal neurons, and astrocyte progenitors from hPSCs

The generation of cortical neural progenitors was performed as previously described (Yan et al., 2016; Yan et al., 2018). Briefly, hPSCs were seeded into Ultra-Low Attachment (ULA) 24-well plates (Corning, Inc., Corning, NY) at 3.0 – 3.5 ×10^5^ cells per well in 1mL of TeSRE8 medium and grown for 2 days. ROCK inhibitor Y27632 (10 µM) was added during the seeding and removed after 24 h. Then, the culture was switched to the neural differentiation medium composed of Dulbecco’s modified Eagle’s medium/nutrient mixture F-12 (DMEM/F12) plus 2% B27 w/o vitamin A serum-free supplement. At day 1 in the neural differentiation medium, the cells were treated with dual SMAD signaling inhibitors: 10 µM SB431542 and 100nM LDN193189 (Sigma). After 7 days, the cells were incubated with cyclopamine (1 µM, Stemgent), FGF-2 (10 ng/mL, R&D System) and epidermal growth factor (EGF, 10 ng/mL, R&D System) for another 8 days. The cell cultures were maintained in FGF-2 until day 21 in suspension and were dissociated by accutase for printing.

The differentiation of GABAergic interneurons was based on our developed protocol(Liu et al., 2013a; Liu et al., 2013b). After 7 days of the above neural differentiation, the sonic hedgehog (SHH) activator purmorphamine (1 µM, Tocris) was added. At day 21, the GABAergic interneuron progenitors were also dissociated to single cells for printing.

The striatal DARPP32^+^ neurons or medium spiny neurons were generated using our previous protocol (Ma et al., 2012). Briefly, 40 ng/mL SHH C25II (R&D Systems) was added to the neural differentiation medium from day 7 to day 25 after one week of dual SMAD signaling inhibitors treatment. At day 25, the striatal progenitors were then treated with VPA (10μM, Sigma) for 5 days. And single striatal neurons were prepared for printing.

The generation of astrocyte progenitors was performed from our previous protocol(Krencik et al., 2011; Li et al., 2018). Briefly, hPSCs were treated with dual SMAD inhibitors to generate neuroepithelia for 14 days on a monolayer culture. On day 14, neuroepithelia were lifted after brief treatment with 500 µM EDTA (Life Technologies), and resuspended DMEM/F12 plus 1% N2 supplement and FGF-2 (10 ng/mL, R&D System). Half medium change was performed every two to three days. On day 30, EGF (10 ng/mL, R&D System) was added to encourage the expansion of glial progenitors till day 180. And single astrocyte progenitors were prepared for printing.

#### Preparation of fibrin gel

Stock solution of 50 mg/mL fibrinogen, 100 U thrombin, 250 mM CaCl_2_ (100X) and 10 mg/mL aprotinin (20X) were prepared in the following manner: Fibrinogen (F3879, Sigma) was dissolved in Dulbecco’s phosphate buffered saline (DPBS) without calcium and magnesium for 4 h at 37 ℃. The solution was sterile-filtered and stored at -80 ℃ for use. CaCl_2_ (Sigma) was dissolved in deionized (DI) water and filtered for use. Thrombin (T7009, Signa) was dissolved in DPBS and sterile-filtered. The solution was stored at -20 ℃ until use. Aprotinin (A1153, Sigma) was dissolved in DPBS and stored at -20 ℃ until use.

#### Culturing neural cells in fibrin gel

Dissociated hPSC-NPCs at a density of 1 × 10^6^ were mixed with fibrinogen at the following concentrations (1, 2.5, 5, 10 and 20 mg/mL). 0.5 mg/mL aprotinin was added for preventing gel degradation. 1 µL of the fibrinogen-laden cells were dropped on the poly-ornithine coated plates at RT (room temperature). And 1 µL of different concentrations of thrombin (0.5, 1, 25,5 and 10 U) solution with 2.5 mM CaCl_2_ were then added and mixed with the cell-fibrinogen solution. Fresh medium was added after gelation. Cells were encapsulated in the fibrin gel constructs and cultured at 37 ℃ incubator for characterizations.

#### Gelation testing of fibrin gel

The gelation testing of the fibrin gel was modified from previously reports (Kubota et al., 2004; Sproul et al., 2018). Different concentrations of fibrinogen solutions and thrombin solutions were prepared. 1 µL fibrinogen solution was dropped onto cover slide and was mixed well with the same volume of thrombin. To test the gelation, a pipette tip of 2 µL was used to touch the gel to check whether the gel was solid or not (no liquid could be taken by the pipette tip). And the time from the addition of thrombin and formation of the gel was recorded. We defined it the gelation time.

#### Bioink preparation

The four groups of hydrogel mixtures (Supplementary Table 1) were used in this study for cell-laden bioinks: gelatin/alginate/fibrinogen (GLN + ALG + FN), alginate/nanofibrillate cellulose/fibrinogen (ALG + NFN + FN), Matrigel/fibrinogen (MG + FN), and hyaluronic acid/fibrinogen (HA + FN). The crosslinking solution was CaCl_2_ + thrombin + transglutaminase (TG). All the bioinks were based on the formation of fibrin gel. The final concentrations of fibrinogen and thrombin in the fibrin gel constructs were maintained at 2. 5 mg/mL and 0.5 U, respectively. All the cells used for printing were disaggregated and were filtered using 30 µm CellTrics^TM^ disposable filter (04-004-2326, Sysmex)to avoid nozzle clogging. Cells were counted through hemocytometer and were prepared at a density of 1 × 10^7^/ mL.

Crosslinking buffer CaCl_2_ + thrombin + transglutaminase (TG) was prepared as following: The TG solution was created as a previously published protocol (Joung et al., 2018; Kolesky et al., 2016). Lyophilized Moo Glue powder (Modernist Pantry; ME, USA) was prepared in DPBS and stirring at 37 ℃ until completely dissolved to make 60 mg/mL solution. The final working crosslinking solution contained 2.5 mM CaCl_2_, 1U thrombin and 0.2 % (w/v) of TG. The crosslinking was performed at 3-5 min at room temperature (RT). To avoid dehydration of gel, the crosslinking agent was added immediately after printing. The volume of the crosslinking solution was equal to the volume of bioinks.

The GLN + ALG + FN was hydrogel mixture of gelatin (GLN), alginate (ALG) and fibrinogen (FN) at a ratio of 2: 1 :1 by volume. Gelatin (G6411, Sigma) was dissolved in DPBS for 12 h at 90 ℃ to make 120 mg/mL stock solution. The solution was stored at 4 ℃ until use. 3% (w/v) of low viscosity alginate (A18565, Alfa Aesar) was prepared in DPBS without calcium and magnesium. The fibrinogen stock (50 mg/mL) was diluted by fresh neural differentiation medium to make 20 mg/mL for use. For printing test, hPSC-NPCs at day 21 were dissociated by accutase and filtered to make single cells. The cells were centrifuged and resuspended with 20 mg/mL fibrinogen solution at a density of 1 × 10^7^/mL. The cell-fibrinogen solution was then mixed with 120 mg/mL gelation and 3% (w/v) alginate solution at a volume ratio of 2: 1: 1. The bioink was prepared freshly before printing. The printing was performed at 10 ℃. After printing, the hydrogel was crosslinked with a thrombin + transglutaminase (TG) + CaCl_2_ for 10-15 min at 10 ℃. And fresh medium was added after crosslinking for post-printing culture at 37 ℃ for characterizations.

The ALG + NFN + FN hydrogel mixture, created from 3% (w/v) alginate solution (ALG), 10 % (w/v) nanofibrillate cellulose (NFN) and 15 mg/mL fibrinogen with a volume ratio of 1: 1: 1. 10 % (w/v) NFN (methyl cellulose, viscosity 15 cPs, 45490, Alfa Aesar), was dissolved in DPBS. The printing was performed at RT. After printing, the hydrogel was crosslinked with a thrombin + transglutaminase (TG) + CaCl_2_ for 3-5 min at RT. And fresh medium was added after crosslinking at 37 ℃ for characterizations.

The MG + FN bioink was prepared from 10 mg/mL fibrinogen solution and Matrigel (BD Biosciences) with a volume ratio of 1: 1. Briefly, cells were centrifuged and resuspended with 10 mg/mL fibrinogen solution at a density of 1 × 10^7^/mL. And Matrigel was added to make the final cell-laden bioink. The printing and crosslinking were performed at RT.

The HA + FN bioink was made of hyaluronic acid (HA) (53747, Sigma) and 10 mg/mL fibrinogen with a volume ratio of 1: 2. 3% (w/v) HA was prepared in DPBS at 37 ℃ and stirred until completely dissolved. The printing was performed at RT. After printing, the hydrogel was crosslinked with a thrombin + transglutaminase (TG) + CaCl_2_ for 3-5 min at RT. And fresh medium was added after crosslinking at 37 ℃ for characterizations.

#### Cell viability assay

Cell viability was indicated by the percent of live cells to total cells from Live/Dead assay. Live/Dead^®^ staining kit (Molecular Probes) was used to assess cell viability. Immediately after harvesting, the cells were incubated in DMEM/F12 containing 1 µM calcein AM and 2 µM ethidium homodimer I for 30 min. Cells were washed and representative images were taken. The number of live (green) and dead (red) cells were counted in each field using Cell Counter in Image-J. The live/dead cell numbers from the five images of one sample were averaged to give each data point and ten samples from three different batches were used to determine the viability.

#### Printing procedure

The printers were from CELLINK^®^, INKREDIBLE+^TM^ and BioX^TM^ (https://www.cellink.com/bioprinting/bio-x/). The printers had two or three printheads, which could deposit multiple types of cells at the same time. The dispensing apparatuses were connected to the printer to extrude different bioinks through the micronozzle with different inner diameters. Before printing, the UV light was turned on for 5 min to make sterile environment for printing. The printing seed, printing pressure and sizes of nozzle were optimized by measurement of the width of printed tissue layer. For printing structure test, we used CELLINK^®^ Start bioink without cells. The micronozzle was 27G (NZ3270005001, Cellink) with a 200 µm inner diameter. The printing speed for the Start bioink was at 3 mm/s (line-dispensing printing mode) and printing pressure was 50 kPa. For the printing with cell-laden bioinks, we performed a printing speed of 5 mm/s (line-dispensing printing mode) and printing pressure of 100 kPa using the blunt needle of 30G (NZ5300505001, Cellink, inner diameter 150 µm) or blunt needle of 27G (NZ6270505001, Cellink, inner diameter 200 µm). All the print paths were controlled using G-code commands, which were generated by the software Slic3R from 3 models.

#### Printing neural tissues

For printing the layered tissue using colored cells (GFP- and mCherry-labeled) or unlabeled cells, neural cells (glutamate neuron progenitors, GABA neuron progenitors, striatal neuron progenitors and astrocyte progenitors) were dissociated and filtered to make single cells. They were prepared separately and laden with the HA + FN bioink at a density of 1 × 10^7^/mL. The gel-laden cells were delivered to two different nozzles and deposited onto poly-ornithine coated coverslips. The crosslinking solution was added immediately after printing. The gelation was performed at RT. Printed tissues were incubated at 37 ℃ in the fresh neural differentiation medium for the following experiments. At day 0 post-printing, printed tissues were cultured in neural basal medium plus 2% B-27 serum-free supplement, 10 ng/mL BDNF (brain-derived neurotrophic factor), 10 ng/mL GDNF (glial cell line derived neurotrophic factor), 200 μM AA (ascorbic acid), and 1 μM cAMP. For the printed tissue with astrocytes, 10 ng/mL CNTF (ciliary neurotrophic factor) was added as well. ROCK inhibitor Y27632 (10 µM) was added for the first 24 h. The medium was changed every 3 days.

#### Immunofluorescence and quantification

Cells on coverslips or wells were rinsed with PBS and fixed in 4% paraformaldehyde for 20 min. After rinsing with PBS twice, cells were treated with 0.3% Triton for 10 min followed by 10% donkey serum for 1 hour before incubating with primary antibodies overnight at 4 ℃. Cells were then incubated for 1h at room temperature with secondary antibodies. The nuclei were stained with Hoechst 33342 (Hoe) (Sigma-Aldrich). Please see key resources table for list of antibodies. Images were taken with a Nikon A1R-Si laser-scanning confocal microscope (Nikon, Tokyo, Japan). The primary antibodies used were listed in Tables S2. Cell quantification was previously described (Krencik et al., 2011). Briefly, multiple fields were chosen randomly under the fluorescent filter for nuclear staining throughout the coverslips in areas which contained a similar density of Hoe^+^ cells and the total cells were counted. The fluorescent filters were shifted during imaging to count the cells labeled by different antibodies in the same field in the same manner. The quantitative data were repeated for three biological replicates.

#### Synaptic puncta quantification

The synaptic puncta analysis was performed as previously described (Cheng et al., 2016; Goldman et al., 2013). Briefly, synapses were identified by the co-localization of the pre- and post-synaptic puncta and 100 µm of neurite proximal to the cell soma was selected for analysis using ImageJ. Images were thresholded using a constant value for each channel to remove low frequency background, and an image generated using the co-localization highlighter to identify regions of overlap between the labeling. Puncta were analyzed in each individual channel and in the co-localization image. About 30 neurons were collected for each condition, and each experiment was repeated for three biological replicates.

#### Lenti-GCaMP6 virus preparation

GCaMP6 lentiviral particles were produced in HEK293FT cells with a 2nd generation lentiviral system. Transfection and lentiviral collection were conducted by Lipofectamine 3000 reagent for lentiviral production. In brief, 7 ×10^6^ HEK293FT cells were seeded on a 10cm plate the day prior to transfection. Next day, the cells were transfected with 4.3 μg of GCaMP6 transfer vector along with 10 μg of the packaging plasmid psPax2 and 3 μg the envelope plasmid pDM2.g (psPax2 [Addgene plasmid #12260] and pDM2.g [Addgene plasmid #12259] were gifts from Didier Tronto). Supernatants containing lentiviral particles were collected 24 hours and 52 hours post-transfection. The success of lentiviral production was confirmed using Lenti-X GoStix (TakaRa Bio) following manufacturer’s instructions. The supernatant was then combined and concentrated with Lenti-X Concentrator (TakaRa Bio). Concentrated lentiviral particles were resuspended in DPBS with 0.1%BSA, aliquoted and stored at -80°C.

#### Lenti-jRGECO1b and lenti-iGluSnFR virus preparation

Lenti-EF1-jRGECO1b and lenti-EF1-iGluSnFR were cloned using lenti-CMV-GFP vector as a backbone(Gao et al., 2020) and the CMV-GFP cassette was replaced with EF1-jRGECO1b and EF1-iGluSnFR, respectively. The jRGECO1b was cloned from pAAV.Syn.NES-jRGECO1b.WPRE.SV40 vector (addgene plasmid #100857)(Marvin et al., 2013). And the iGluSnFR was cloned from pENN.AAV.GFAP.iGluSnFr.WPRE.SV40 vector (addgene plasmid #98930). The sequence of the inserted cassette was confirmed by sequencing. Lentivirus production was performed as described previously (Gao et al., 2020) with modifications. Briefly, the viral transfer vector DNA and packaging plasmid DNA were co-transfected into HEK293T cells using PEI. The medium containing lentivirus was collected at 36, 60 and 84 h post-transfection, pooled, filtered through a 0.2-μm filter, and concentrated using an ultracentrifuge at 19,400 rpm for 2 h at 4°C using a SW32Ti rotor (Beckman). The virus was washed once and then resuspended in 50 μl PBS. We routinely obtained 5 ×10^8^ infectious viral particles /ml for lentivirus.

#### Calcium and Glutamate imaging

Human PSC-derived astrocyte progenitors at day 180 were infected with Lenti-GCaMP6 or Lenti-iGluSnFR. After 3 days, the virus infected astrocyte progenitors were washed and then printed with hPSC-derived cortical and MGE progenitors at ratio of 4: 5: 1. Calcium or glutamate imaging was performed as previously described (Sloan et al., 2017). Briefly, the printed tissues were washed with low potassium Tyrode’s solution (Low-KCl) (2 mM KCl, 129 mM NaCl, 2 mM CaCl_2_, 1 mM MgCl_2_, 30 mM glucose, 25 mM HEPES, 0.1% and Bovine Serum Albumin, pH 7.4) three times and incubated with the solution for 30 min at 37 ℃. When imaging, the sample was placed on the stage of confocal fluorescence microscope (A1, Nikon). A high potassium Tyrode’s solution (High-KCl) (67 mM KCl, 67 mM NaCl, 2 mM CaCl_2_, 1 mM MgCl_2_, 30 mM glucose, 25 mM HEPES, 0.1% and Bovine Serum Albumin, pH 7.4) was then applied. As a control, the virus infected astrocyte progenitors were also cultured alone for the same period and High-KCl solution was applied. ImageJ was used for the following analysis. The fluorescence change was defined as ΔF/F(t) = (F_0_-F(t))/F_0_, where F_0_ is the average fluorescence intensity of the imaging area for samples in the Low-KCl solution, F(t) was the fluorescence intensity at a given time. The ΔF/F of the printed tissue was normalized by comparison of the ΔF/F of the control.

#### Optogenetic stimulation

Human PSC-derived striatal neuron progenitors at day 21 were infected with Lenti-jRGECO1b. After 3 days, the virus infected striatal progenitors were then printed with hPSC-ChR2-EYFP derived cortical progenitors. Optogenetic stimulation was performed as previously described (Dong et al., 2020). Briefly, the printed cortical and striatal tissues were placed in a 35-mm glass-botton dish in neural medium and imaged using confocal fluorescence microscope (A1, Nikon). For optogenetic stimulation, ChR2-cells were activated with 470 nm light using a custom-made LED device (1 Watts, 470 nm; Cree lighting Inc.) coupled to a fiber optic cable. jRGECO1b was imaged at a rate of 4.6 frames per second. Stimulation experiments included 1450 frames, and 470 nm LED light was applied every 200 - 300 frames. ImageJ was used for the Ca^2+^ wave analysis.

#### Electrophysiology

Whole-cell patch-clamp recordings were made from human PSC-derived cortical glutamatergic and GABAergic neurons. The bath solution consisted of 135 mM NaCl, 3 mM KCl, 2 mM CaCl2, 1 mM MgCl2, 10 mM HEPES, 11 mM glucose, 10 mM sucrose, pH7.4. Recording pipettes were filled with an intracellular solution containing 120 mM potassium D-gluconate, 1 mM ethylene glycol-bis (β-aminoethyl ether) N,N,N’,N’-tetraacetic acid (EGTA), 10 mM 4-(2-hydroxyethyl)piperazine-1-ethanesulfonic acid (HEPES), 4 mM ATP-Mg, 0.3 mM GTP-Na, 10 mM phosphocreatine, 0.1 mM CaCl2, 1 mM MgCl2, pH 7.2, 280–290 mOsm/L. Printed tissues on the coverslips were transferred into a perfusion chamber. Individual cells were visualized with the help of an infrared differential interference contrast (IR-DIC) Olympus BX51WI microscope at 40× water-immersion objective, and different cell types were further identified by their fluorescence detected with a CCD camera and displayed on a monitor. Recording pipettes were made by pulling the glass (BF150-86-10, Sutter Instrument) onto a P-97 Flaming/Brown micropipette puller (Sutter Instrument). The pipette resistance was typically 3–6 MΩ after being filled with the intracellular solution. Since there were multiple layers of cells, we placed the pipette tip into the bath solution and focused on the tip. Once the pipette tip touched the target cells and formed a very small dimple, we released the positive pressure to obtain a GΩ seal. We then gave a small negative pressure to break through the membrane before recording according to our protocol. Briefly, the neurons were held at −70 mV to record the Na+/K+ channel activities with the voltage-clamp model. For recording action potentials, the cells were held at 0 pA with the current-clamp model, and with the steps of injected currents from −50 pA to + 50 pA. Spontaneous excitatory postsynaptic currents (sEPSCs) and spontaneous inhibitory postsynaptic currents (sIPSCs) were recorded in gap-free mode at a holding potential of -70 mV and 0 mV, respectively. sIPSCs were blocked by application of the GABAA receptor antagonist bicuculline (10 µM). For ChR2 stimulation, a light stimulation fiber was placed 5mm from the dish. A custom-made LED device (1 Watt, 470 nm; Cree lighting Inc.) coupled to a fiber optic cable was used to achieve the stimulation. EPSCs were recorded before the blue light stimulation(baseline), with blue light stimulation (30∼60 s, 470 nm, 0.4 mW/mm2) and after blue light stimulation under the voltage-clamp mode. mCherry-striatal neurons in printed cortico-striatal tissue were randomly selected for recordings. An Olympus BX51WI microscope was used to visualize neurons. A MultiClamp 700B amplifier (Axon instruments, Molecular Devices, Sunnyvale, CA, USA) was used to investigate the voltage clamp and current clamp recordings. Signals were filtered at 4 kHZ using a Digidata 1550B analog-digital converter (Axon instruments) and stored for further analysis. Data were analyzed with Clampfit 11.0.3 (Axon instruments), GraphPad Prism 5 (GraphPad Software Inc., La Jolla, CA, USA), CorelDraw 2019 (Corel, Canada), Igor 4.0 (WaveMetrics, Lake Oswego, OR, USA).

#### Statistics

All data were expressed as mean ± standard deviation. Graphs and statistical analysis were made in GraphPad Prism. Distribution of the raw data was tested for normality of distribution; statistical analyses were performed using the t-test, or ANOVA tests as indicated.

**Table S1.** Comparison of 3D bioprinted human neural tissues capturing CNS functions

