## Supplemental figures for "3D Bioprinting of Human Neural Tissues with Functional Connectivity"

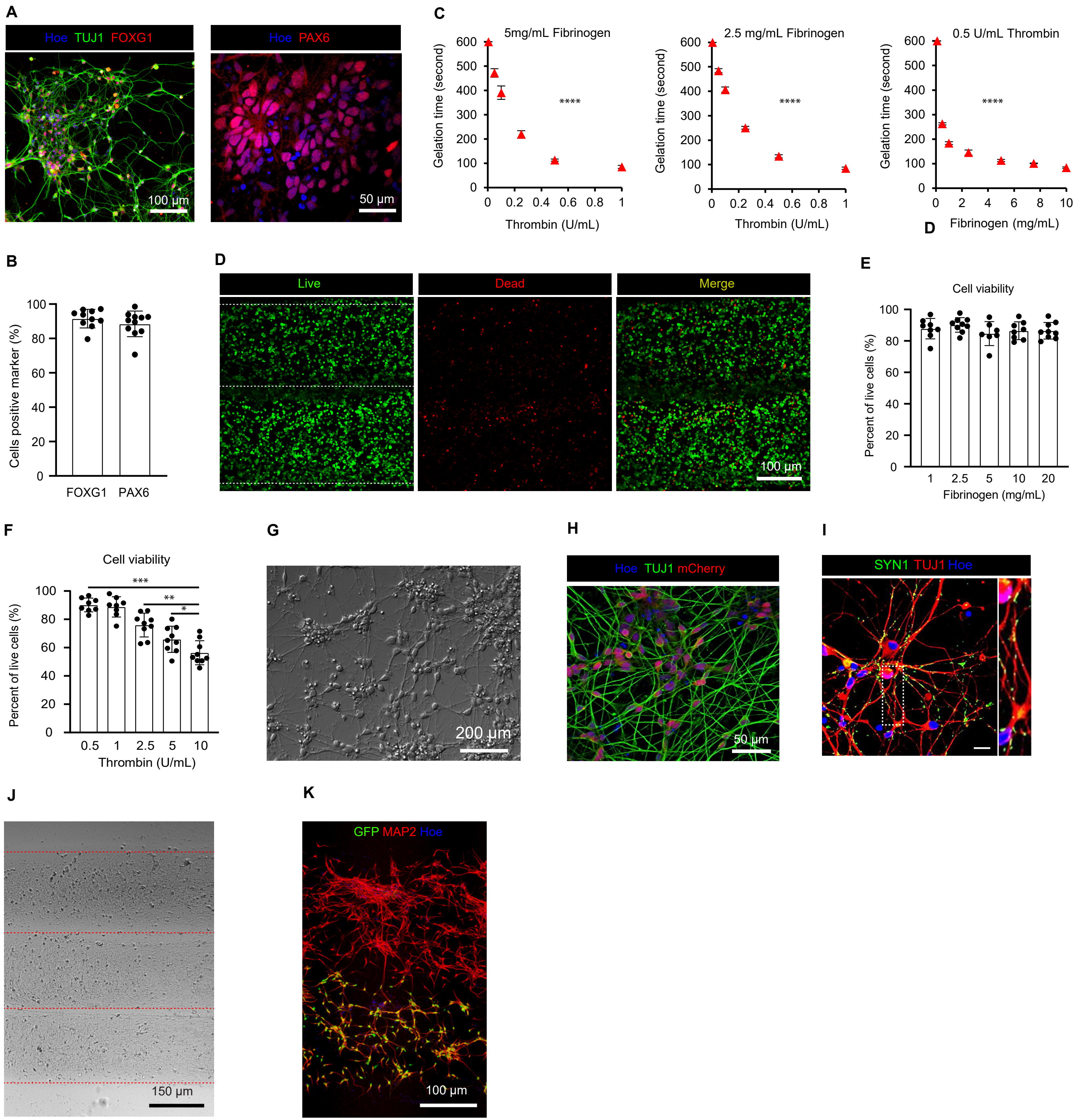
A
Hoe TUJ1 FOXG1
100 µm
Hoe PAX6
50 µm
C
5mg/mL Fibrinogen
Gelation time (second)
Thrombin (U/mL)
****
2.5 mg/mL Fibrinogen
Gelation time (second)
Thrombin (U/mL)
****
0.5 U/mL Thrombin
Gelation time (second)
Fibrinogen (mg/mL)
****
D
B
Cells positive marker (%)
FOXG1 PAX6
D
Live
Dead
Merge
100 µm
E
Cell viability
Percent of live cells (%)
1 2.5 5 10 20
Fibrinogen (mg/mL)
F
Cell viability
Percent of live cells (%)
***
**
*
0.5 1 2.5 5 10
Thrombin (U/mL)
G
200 µm
H
Hoe TUJ1 mCherry
50 µm
I
SYN1 TUJ1 Hoe
J
150 µm
K
GFP MAP2 Hoe
100 µm

**Figure S1. Characterization of human neural cell growth in the fibrin gel, Related to Figure 1 and Figure 2**

(A) Immunostaining of hPSC-derived cortical NPCs for neural markers FOXG1 and PAX6.

(B) Quantitative analysis of hPSC-derived cortical NPCs for the expression of FOXG1 and PAX6 (n = 10-11 from three different experiments).

(C) Gelation testing of fibrin gel composition. (5 mg/mL Fibrinogen: One-way ANOVA; interaction $F(5,30) = 2112$; ****$P < 0.0001$; 2.5 mg/mL Fibrinogen: One-way ANOVA; interaction $F(5,12) = 2629$; ****$P < 0.0001$; 0.5 U/mL Thrombin: One-way ANOVA; interaction $F(6, 14) = 3679$; ****$P < 0.0001$; 6 smaples for each testing point from three independent experiments.)

(D) Live/Dead assay of two-layered cells using HA + FN as a bioink for printing.

(E) and (F) The viability (percentage of live cells) of neural cells under different concentrations of fibrinogen and thrombin (one-way ANOVA; interaction $F(4, 38) = 29.25$, ***$P < 0.001$; one-way ANOVA; interaction $F(2, 24) = 11.32$, **$P < 0.01$; unpaired t-test; *$P < 0.05$; 8-10 samples from three different experiments).

(G) Bright phase image of NPCs grown in the fibrin gel for one month.

(H) Immunostaining for neural markers TUJ1 and mCherry at day 4 in the gel.

(I) Immunostaining for synaptic markers SYN1 with TUJ1 at day 30 in the fibrin gel.

(J) Phase contrast images of printed layers at day 2 after printing.

(K) Immunostaining for $GFP^{+}$ layer and $GFP^{-}$ layer with MAP2 staining at day 7 after printing.

Data are represented as mean ± SEM. Hoe, Hoechst 33342.

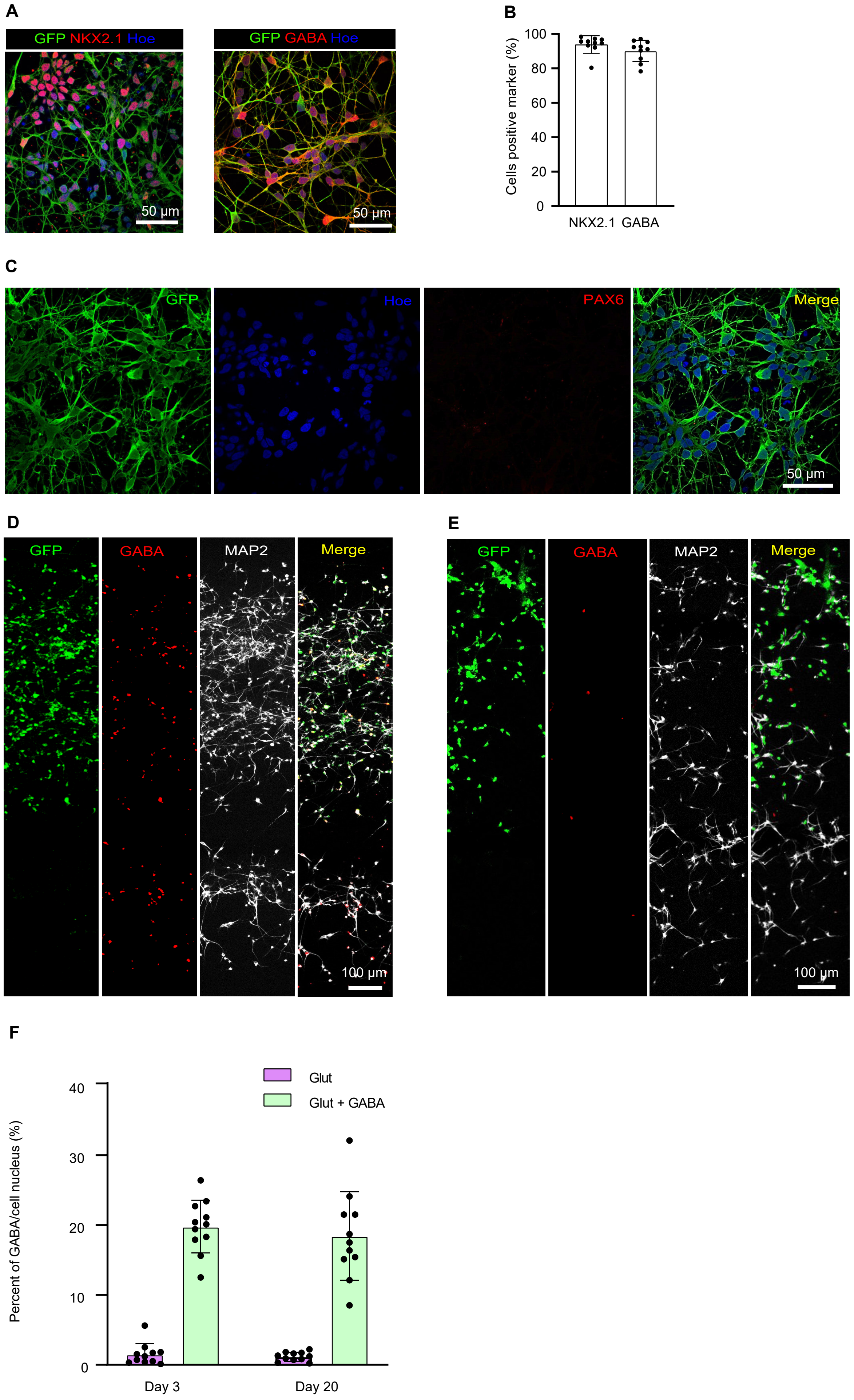

A
GFP NKX2.1 Hoe
50 μm
GFP GABA Hoe
50 μm
B
Cells positive marker (%)
100
80
60
40
20
0
NKX2.1
GABA
C
GFP
Hoe
PAX6
Merge
50 μm
D
GFP
GABA
MAP2
Merge
100 μm
E
GFP
GABA
MAP2
Merge
100 μm
F
Glut
Glut + GABA
Percent of GABA/cell nucleus (%)
40
30
20
10
0
Day 3
Day 20

**Figure S2. Printing tissues with cortical and MGE progenitors, Related to Figure 3**

(A) Immunostaining of hPSC-derived MGE progenitors for GFP and NKX2.1 and GABA.

(B) Quantification of NKX2.1 and GABA populations (10 samples from three different batches).

(C) Immunostaining of hPSC-derived MGE progenitors for GFP and PAX6, showing negative expression of PAX6.

(D) Immunostaining of printed tissues (with cortical and MGE progenitors) for GABA and MAP2 at day 20 after printing.

(E) Immunostaining of printed tissues (with cortical progenitors-only) for GABA and MAP2 at day 3 after printing.

(F) Quantitative analysis of GABA expression for different groups at day 3 and 20 after printing (11 samples from three different batches for each condition).

Data are represented as mean ± SEM. Hoe: Hoechst 33342.

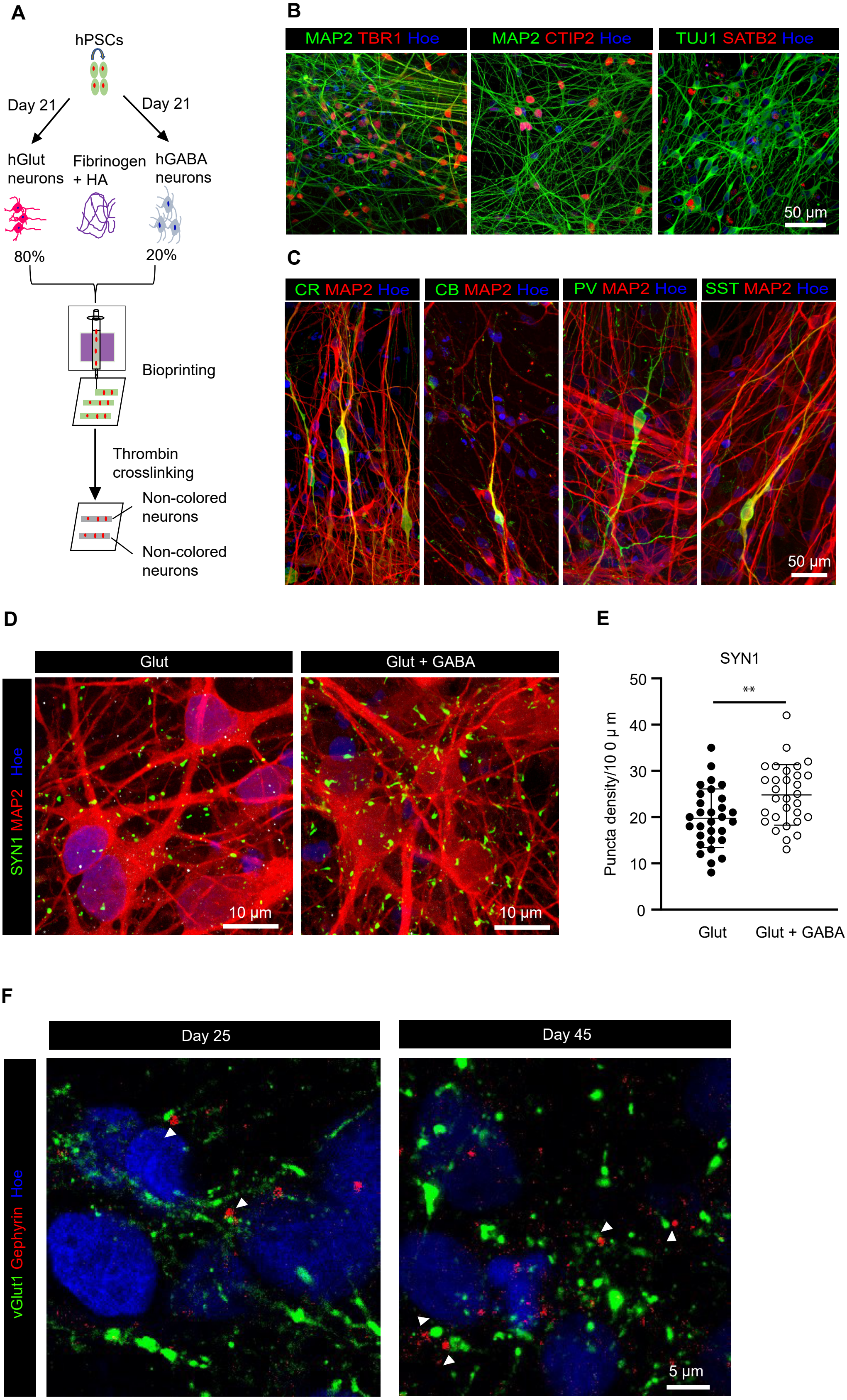
A
hPSCs
Day 21
Day 21
hGlut neurons
Fibrinogen + HA
hGABA neurons
80%
20%
Bioprinting
Thrombin crosslinking
Non-colored neurons
Non-colored neurons
B
MAP2 TBR1 Hoe
MAP2 CTIP2 Hoe
TUJ1 SATB2 Hoe
50 μm
C
CR MAP2 Hoe
CB MAP2 Hoe
PV MAP2 Hoe
SST MAP2 Hoe
50 μm
D
Glut
Glut + GABA
SYN1 MAP2 Hoe
10 μm
10 μm
E
SYN1
**
Puncta density/10 0 μ m
0
10
20
30
40
50
Glut
Glut + GABA
F
Day 25
Day 45
vGlut1 Gephyrin Hoe
5 μm

**Figure S3. Characterization of the tissues printed with cortical and GABAergic neurons, Related to Figure 3**

(A) Design of the experiments.

(B) Immunostaining of printed tissues for cortical neuron markers TBR1, CTIP2 and SATB2 at 60 days after printing.

(C) Immunostaining of printed tissues for markers of GABAergic neuronal subtypes calretinin (CR), calbindin (CB), parvalbumin (PV) and somatostatin (SST) with MAP2 at day 60 after printing.

(D) Immunostaining of different printed tissues for SYN1 puncta at day 20 after printing.

(E) Quantitative analysis of SYN1 puncta density for different groups at day 20 after printing (t-test, $^{**}P < 0.01$, 30 neurons from three different batches).

(F) Immunostaining of printed tissues with cortical and MGE progenitors for vGlut1 and Gephyrin puncta at day 25 and 45 after printing. White triagnles indicate the co-localized punctas of vGlut1 and Gephyrin.

Data are represented as mean ± SEM. Hoe: Hoechst 33342.

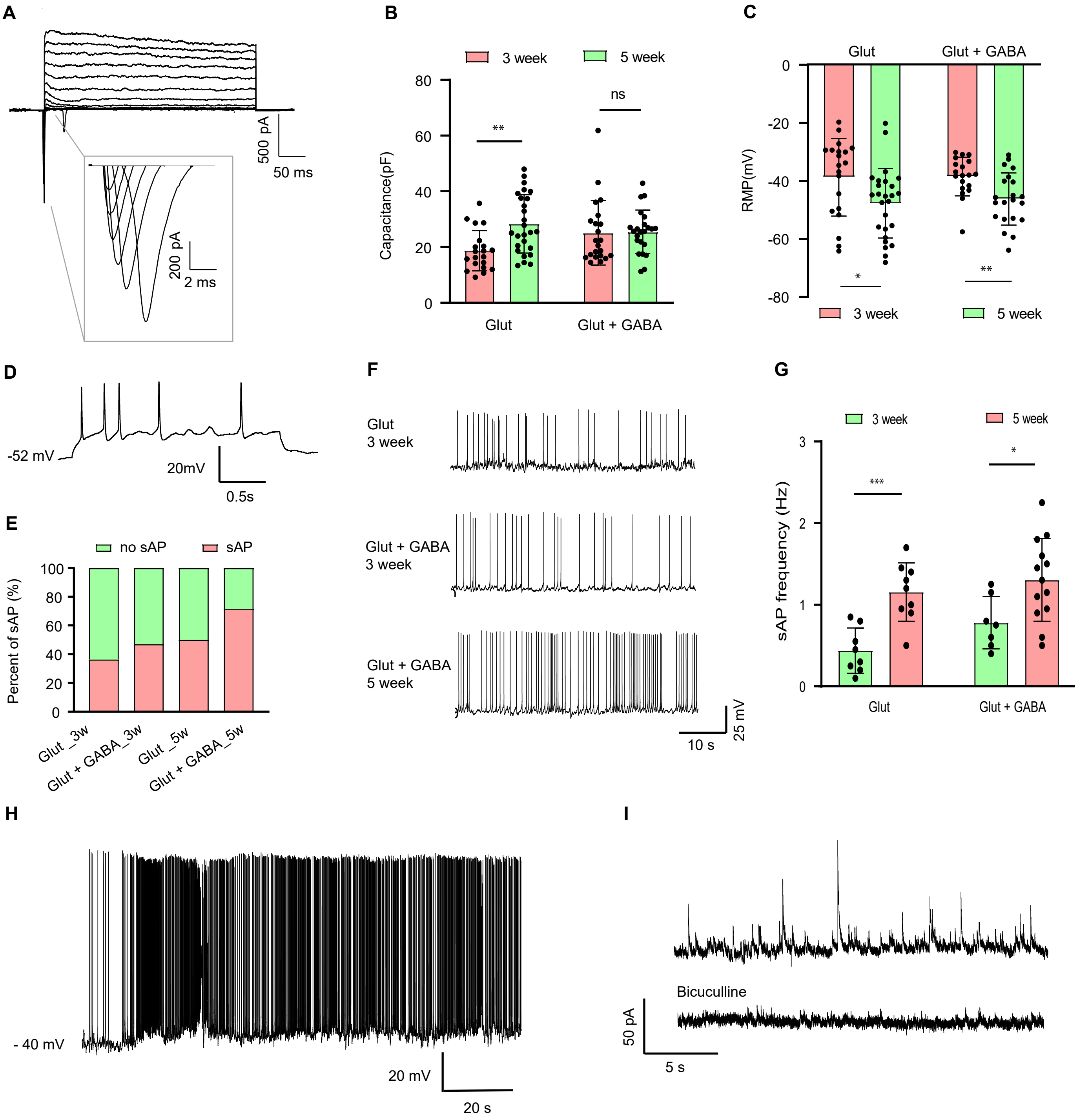
A
500 pA
50 ms
200 pA
2 ms
B
3 week
5 week
80
60
40
20
0
Capacitance(pF)
**
ns
Glut
Glut + GABA
C
Glut
Glut + GABA
0
-20
-40
-60
-80
RMP(mV)
*
**
3 week
5 week
D
-52 mV
20mV
0.5s
E
no sAP
sAP
100
80
60
40
20
0
Percent of sAP (%)
Glut _3w
Glut + GABA_3w
Glut _5w
Glut + GABA_5w
F
Glut
3 week
Glut + GABA
3 week
Glut + GABA
5 week
10 s
25 mV
G
3 week
5 week
3
2
1
0
sAP frequency (Hz)
***
*
Glut
Glut + GABA
H
- 40 mV
20 mV
20 s
I
Bicuculline
50 pA
5 s

**Figure S4. Electrophysiological recordings of printed neural tissues, Related to Figure 3**

(A) The $Na^+/K^+$ current of printed tissues.

(B) The quantitative comparison of capacitance for printed tissue groups at 3- and 5-weeks post-printing (Number of neurons patched: 20 for Glut_3 week, 22 for Glut + GABA_3 week, 26 for Glut_5 week and 23 for Glut + GABA_5 week; multiple t-test, **P < 0.01; P = 0.24).

(C) The quantitative comparison of resting membrane potential (RMP) for printed tissue groups at 3- and 5-weeks post-printing (Number of neurons patched: 22 for Glut_3 week, 17 for Glut + GABA_3 week, 18 for Glut_5 week and 21 for Glut + GABA_5 week; multiple t-test, *P < 0.05; **P < 0.01).

(D) Representative evoked action potential (eAP) in glutamatergic neurons from printed tissues with incorporation of cortical and GABAergic neurons.

(E) The percent of spontaneous action potentials (sAP) for different samples at 3 and 5 weeks (Number of neurons patched: 22 for Glut_3 week, 17 for Glut + GABA_3 week, 18 for Glut_5 week and 21 for Glut + GABA_5 week).

(F) Representative sAP and (G) frequency of sAP from printed tissues at 3 and 5 weeks after printing (Number of neurons patched: 8 for Glut_3 week, 7 for Glut + GABA_3 week, 9 for Glut_5 week and 13 for Glut + GABA_5 week; multiple t-test, ***P < 0.001; *P < 0.05).

(H) Representative sAP of GABAergic neurons from printed tissues with incorporation of cortical and GABAergic neurons.

(I) Representative IPSCs in GABAergic neurons from printed tissues.

Data are represented as mean ± SEM.

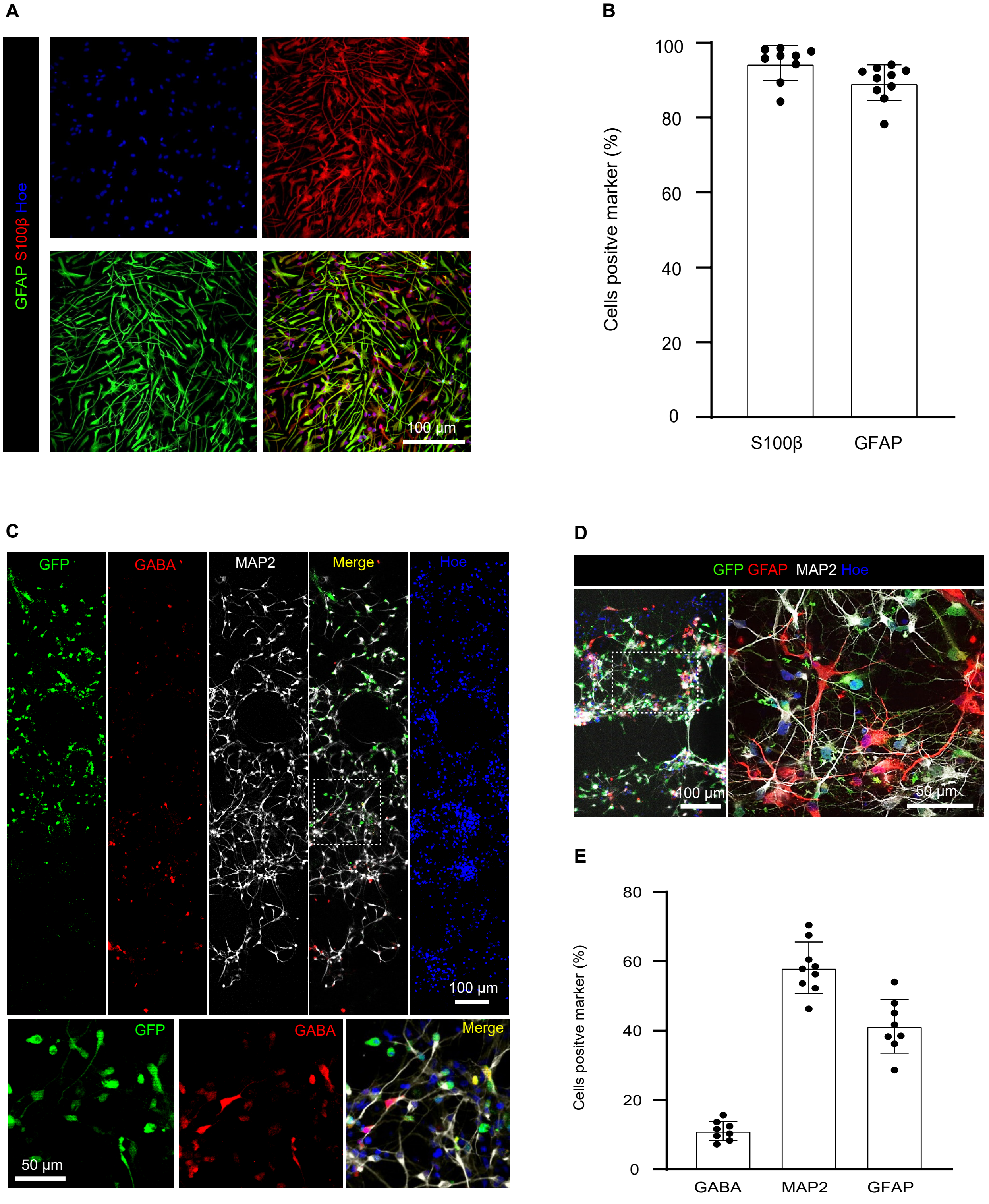

A
GFAP S100β Hoe
100 µm
B
Cells positve marker (%)
100
80
60
40
20
0
S100β
GFAP
C
GFP
GABA
MAP2
Merge
Hoe
100 µm
GFP
GABA
Merge
50 µm
D
GFP GFAP MAP2 Hoe
100 µm
50 µm
E
Cells positve marker (%)
80
60
40
20
0
GABA
MAP2
GFAP

**Figure S5. Printing tissues with neurons and astrocytes, Related to Figure 4**

(A) Immunostaining of hPSC-derived astrocytes for S100β and GFAP.

(B) Quantification of $S100\beta^+$ (9 samples from three different batches) and $GFAP^+$ (10 samples from three different batches) cells among total cells.

(C) Immunostaining of printed tissues with neurons and astrocytes for GABA and MAP2 at day 3 after printing.

(D) Immunostaining of printed tissues with neurons and astrocytes for GFAP and MAP2 at day 13 after printing.

(E) Quantitative analysis of GABA (8 samples from three different batches), MAP2 (9 samples from three different batches) and GFAP (8 samples from three different batches) expression at day 20 after printing.

Data are represented as mean ± SEM. Hoe: Hoechst 33342.

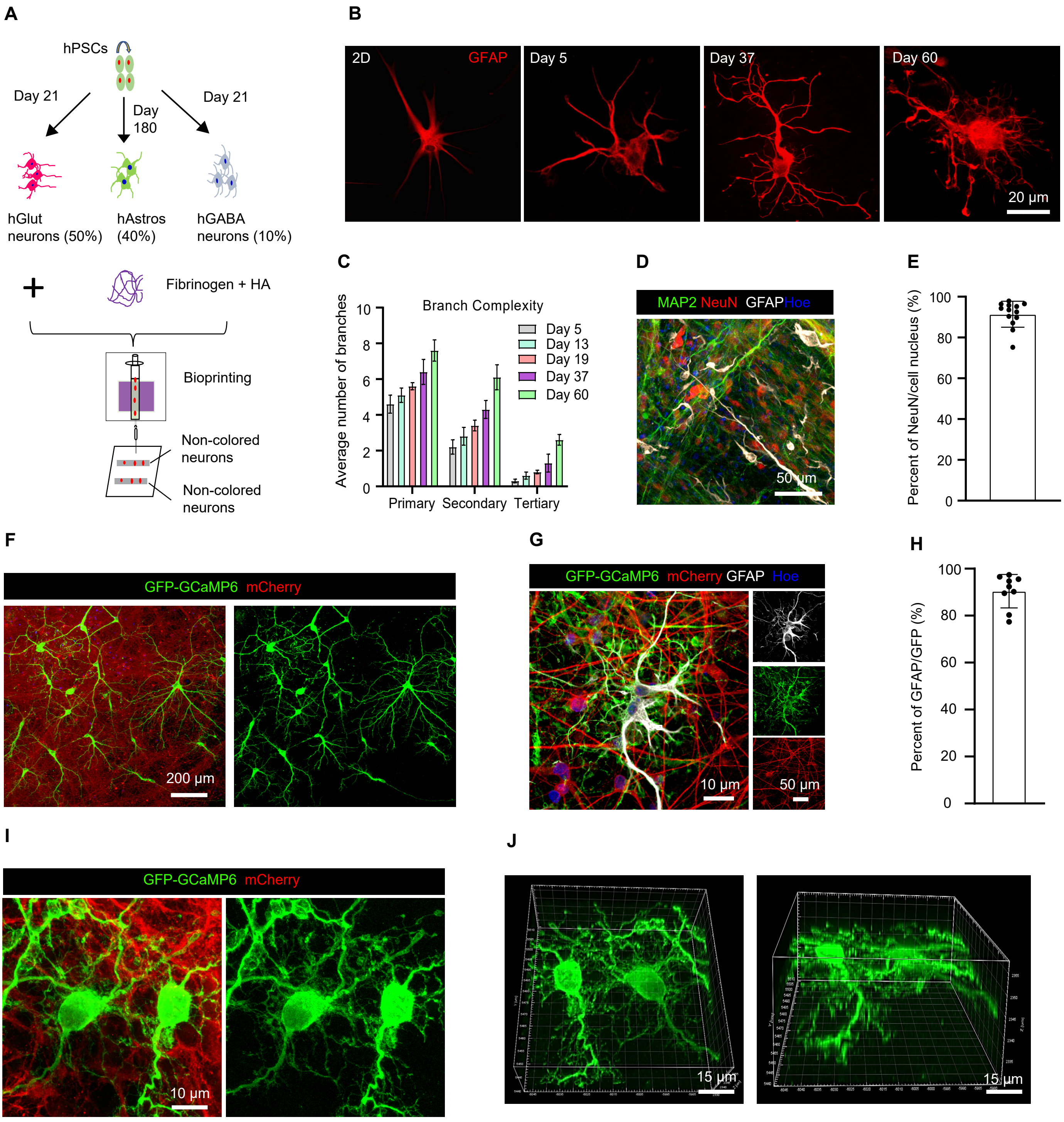
A
hPSCs
Day 21
Day 180
Day 21
hGlut neurons (50%)
hAstros (40%)
hGABA neurons (10%)
+
Fibrinogen + HA
Bioprinting
Non-colored neurons
Non-colored neurons
B
2D
GFAP
Day 5
Day 37
Day 60
20 µm
C
Branch Complexity
Average number of branches
Day 5
Day 13
Day 19
Day 37
Day 60
Primary
Secondary
Tertiary
D
MAP2 NeuN GFAP Hoe
50 µm
E
Percent of NeuN/cell nucleus (%)
F
GFP-GCaMP6 mCherry
200 µm
G
GFP-GCaMP6 mCherry GFAP Hoe
10 µm
50 µm
H
Percent of GFAP/GFP (%)
I
GFP-GCaMP6 mCherry
10 µm
J
15 µm
15 µm

**Figure S6. Characterization of the tissues printed with neurons and astrocytes, Related to Figure 4**

(A) Design of the experiments.

(B) Morphology of astrocytes in 2D cultures and in printed tissues over time.

(C) Quantitative analysis of branch complexity of astrocytes over time.

(D) Immunostaining of printed tissues for GFAP, NeuN and MAP2 at day 30 after printing. (E) Quantitative analysis of NeuN expression (13 samples from three different batches). (F) Immunostaining for GFP-GCaMP6 and mCherry to show the distribution of neurons and astrocytes in printed tissues.

(G) Immunostaining for GFP-GCaMP6 and GFAP at day 30 after printing.

(H) Quantitative analysis of $GFP^+$ and $GFAP^+$ astrocytes (9 samples from three different batches).

(I) Immunostaining for GFP-GCaMP6 and mCherry in printed tissues.

(J) 3D image of (I).

Data are represented as mean ± SEM. Hoe: Hoechst 33342.

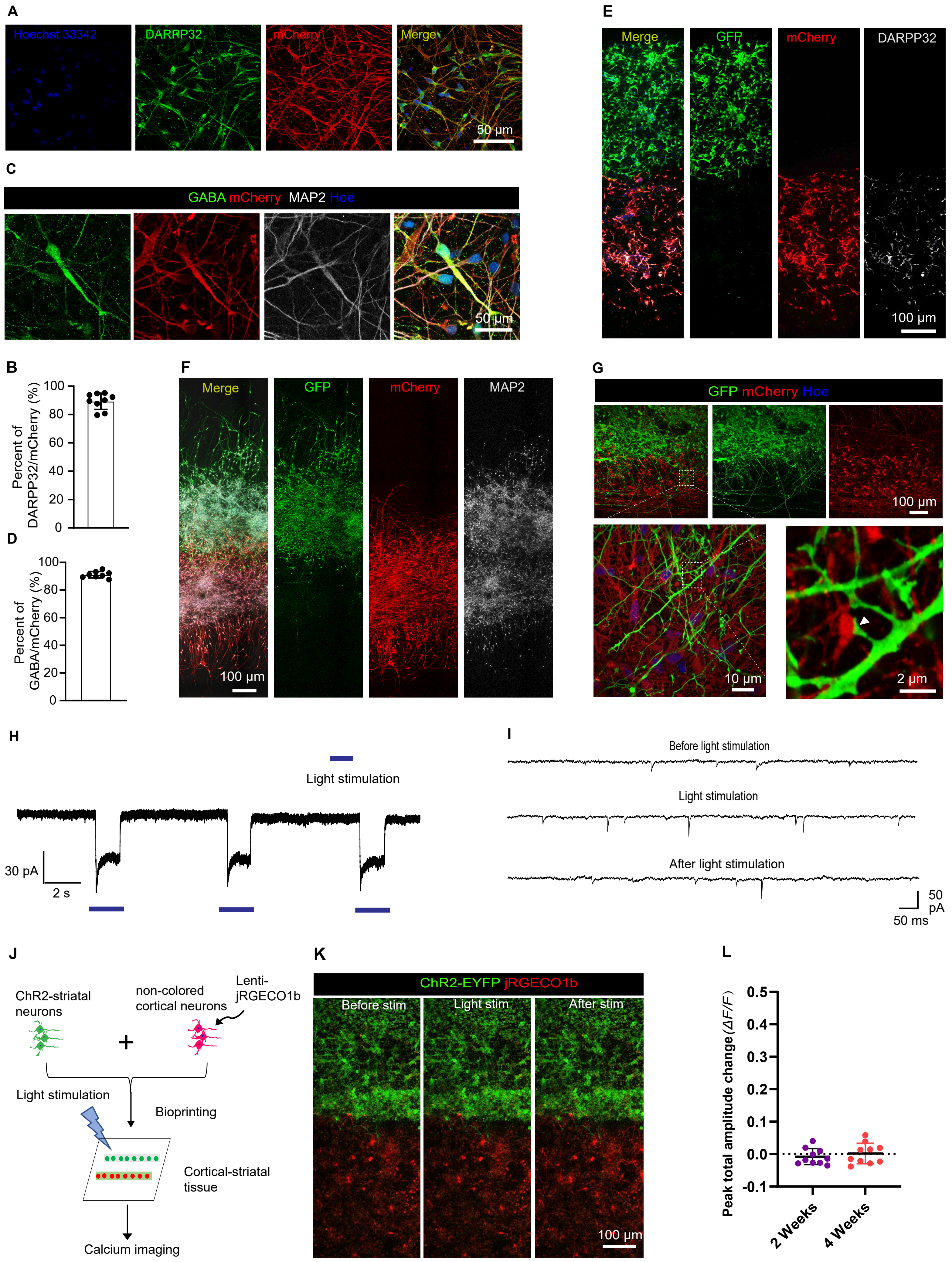
A
Hoechst 33342
DARPP32
mCherry
Merge
50 µm
C
GABA mCherry MAP2 Hoe
50 µm
E
Merge
GFP
mCherry
DARPP32
100 µm
B
Percent of DARPP32/mCherry (%)
100
80
60
40
20
0
D
Percent of GABA/mCherry (%)
100
80
60
40
20
0
F
Merge
GFP
mCherry
MAP2
100 µm
G
GFP mCherry Hoe
100 µm
10 µm
2 µm
H
Light stimulation
30 pA
2 s
I
Before light stimulation
Light stimulation
After light stimulation
50 pA
50 ms
J
ChR2-striatal neurons
non-colored cortical neurons
Lenti-jRGECO1b
+
Light stimulation
Bioprinting
Cortical-striatal tissue
Calcium imaging
K
ChR2-EYFP jRGECO1b
Before stim
Light stim
After stim
100 µm
L
Peak total amplitude change (ΔF/F)
0.5
0.4
0.3
0.2
0.1
0.0
-0.1
2 Weeks
4 Weeks

**Figure S7. Characterization of the printed cortical-striatal tissue, Related to Figure 6**

(A) Immunostaining of the striatal neurons for DARPP32 and mCherry at day 35.

(B) Quantitative analysis of DARPP32$^+$ and mCherry$^+$ neurons (9 samples from three different batches).

(C) Immunostaining of the striatal neurons for GABA and mCherry at day 35.

(D) Quantitative analysis of GABA$^+$ and mCherry$^+$ neurons (8 samples from three different batches).

(E) Representative images of printed cortical-striatal tissues for GFP, mCherry and DARPP32 staining 5 days after printing.

(F) Immunostaining of printed tissues for GFP, mCherry and MAP2 at 2 weeks after printing.

(G) Neurites cross the cortical-striatal layer boundary; arrowhead indicates contact between the GFP+ cortical neurite and mCherry+ striatal neuron at 21 days after printing. See Video S3 shows 3D reconstruction of neuron-neuron interactions.

(H) Induced inward current of ChR2-cortical neurons with light stimulation.

(I) Expanded timescale of spontaneous EPSCs recorded from the striatal neurons within the printed tissues before, during and after the light stimulation 5 weeks after printing.

(J) Schematic diagram illustrating calcium imaging of printed tissues with cortical and striatal neurons. Striatal neurons were ChR2 cells, and cortical neurons were non-colored and infected with lentivirus-jRGECO1b before printing.

(K) Representive calcium images of printed cortical-striatal tissue with light-stimulation of striatal neurons.

(L) Total jRGECO1b amplitudes of $\Delta F/F$ in tissues (2 and 4 weeks post printing) after light-stimulation (10 samples from three different batches).

Data are represented as mean ± SEM. Hoe: Hoechst 33342.

**Table S1.** Comparison of 3D bioprinted human neural tissues capturing CNS functions

**Table S2.** Printing test of the Fibrin gel mixing with other gels

| Hydrogels | Ratio (V) | Crosslink | Printability | Cell viability | Others |
|---|---|---|---|---|---|
| GLN + ALG +FN | 2: 1: 1 | $CaCl_2$+ thrombin + TG | Intermediate | 30.66 ± 4.33% | Printed structure is difficult to preserve |
| ALG + NFC +FN | 1: 1: 1 | $CaCl_2$+ thrombin + TG | High | 5.84 ± 2.67% | |
| MG + FN | 1: 1 | $CaCl_2$+ thrombin + TG | Low | 92.19 ± 4.26% | Nozzle clogging |
| HA + FN | 1: 1 | $CaCl_2$+ thrombin + TG | High | 85.13 ± 6.05% | |

Notes: GLN: gelatin, ALG: alginate, FN: fibrinogen, NFC: nanofibrillated cellulose, MG: Matrigel, HA: hyaluronic acid, TG: transglutaminase, RT: room temperature
