## Supplementary material for "3D Bioprinting of Human Neural Tissues with Functional Connectivity": Resource table

Key Resource Table

| **REAGENT or RESOURCE** | **SOURCE** | | **IDENTIFIER** |
| --- | --- | --- | --- |
| Antibodies |  | |  |
| Rabbit Calbindin | Abcam | | ab25085, RRID: AB_448597 |
| Rabbit Calretinin | Epitomics Inc. | | 2624-1, RRID:AB_2228336 |
| Rat CTIP2 | Abcam | | ab18465, RRID: AB_2064130 |
| Rabbit DARPP32 | Abcam | | Ab40801, RRID: AB_731843 |
| Rabbit DARPP32 | Millipore Sigma | | AB10518, RRID: AB_10807019 |
| Rabbit Drebrin | Millipore | | AB10140, RRID: AB_1977159 |
| Rabbit FOXG1 | Abcam | | ab18259, RRID: AB_732415 |
| Rabbit GABA | Sigma | | A2052, RRID: AB_2314459 |
| Mouse Gephyrin | Synaptic Systems | | 147011, RRID: AB_2810215 |
| Mouse GFAP | Millipore | | IF03L, RRID: AB_2294571 |
| Rabbit GFAP | Dako | | Z033429, RRID:AB_10013382 |
| Mouse GLT1 | BD Transduction Laboratories | | 611654, RRID: AB_399172 |
| Mouse GFP | Millipore | | MAB3580, RRID: AB_94936 |
| Rabbit GFP | Millipore Sigma | | AB3080, RRID: AB_91337 |
| Chicken GFP | Novus Biologicals | | NB100-1614, RRID: AB_10001164 |
| Rabbit MAP2 | Millipore | | AB5622, RRID: AB_91939 |
| Mouse MAP2 | Sigma | | M1406-2ML, RRID: AB_477171 |
| Chicken MAP2 | Abcam | | ab5392, RRID: AB_2138153 |
| Rat mCherry | ThermoFisher Scientific | | M11217, RRID: AB_2536611 |
| Mouse NeuN | Millipore | | MAB377, RRID: AB_2298772 |
| Mouse NKX2.1 | Millipore | | MAB5460, RRID: AB_571072 |
| Goat OTX2 | R&D System | | AF1979, RRID: AB_2157172 |
| Mouse Parvalbumin | Millipore | | MAB1572, RRID: AB_2174013 |
| Mouse PAX6 | DSHB | | PAX6, RRID: AB_528427 |
| Mouse PSD95 | Synaptic Systems | | 124011, RRID: AB_2619799 |
| Mouse S100β | Abcam | | ab11178, RRID: AB_297817 |
| Mouse SATB2 | Abcam | | ab51502, RRID: AB_882455 |
| Mouse SMI312 | BioLegend | | 837904, RRID: AB_2566782 |
| Rat Somatostatin | Millipore | | MAB354, RRID: AB_2255365 |
| Rabbit SOX2 | Millipore | | AB5603, RRID: AB_2286686 |
| Rabbit SYN1 | Synaptic Systems | | 106-003, RRID: AB_2619773 |
| Rabbit TBR1 | Abcam | | ab31940, RRID: AB_2200219 |
| Rabbit TUJ1 | Covance | | PBR-435P, RRID: AB_291637 |
| Mouse TUJ1 | Abcam | | Ab78078, RRID: AB_2256751 |
| Rabbit vGAT | Synaptic Systems | | 131002, RRID: AB_887871 |
| Rabbit vGlut1 | Synaptic Systems | | 135303, RRID: AB_887875 |
| Chemicals and proteins |  | |  |
| Accutase | Innovative Cell Technologies, Inc. | | AT-104 |
| Aprotinin | Sigma | | A1153 |
| B27 supplement w/o Vitamin A | Thermo Fisher Scientific | | 12587-010 |
| Bicuculline methiodide | Tocris | | 2503 |
| Bovine serum albumin (BSA) | VMR | | 10842-772 |
| CaCl_2_ | Sigma | | C7902 |
| c-AMP | Sigma-Aldrich | | D0627 |
| *Continued* |  | |  |
| **REAGENT or RESOURCE** | **SOURCE** | | **IDENTIFIER** |
| Cyclopamine | Stemgent | | 04-0022 |
| D-(+)-Glucose | Sigma | | G8270-1KG |
| Dispase | Thermo Fisher Scientific | | 17105-041 |
| DMEM/F12 basal medium | Thermo Fisher Scientific | | 11330-032 |
| DMH1 | Tocris | | 4126 |
| DPBS | Thermo Fisher Scientific | | 141901144 |
| EGF | R&D Systems | | 236-EG-01M |
| EGTA | Sigma | | E3889-500G |
| FGF-2 | R&D Systems | | 233-FB-500/CF |
| Fibrinogen | Sigma | | F3879 |
| Gelatin | Sigma | | G9391-500G |
| GlutaMAX | Gibco | | 35050-079 |
| GTP-Na | Sigma | | 10106399001 |
| HEPES (1 M) | Thermo Fisher Scientific | | 15630080 |
| Hoechst 33342 | Thermo Fisher Scientific | | R37165 |
| Hyaluronic acid | Sigma | | 53747 |
| KnockOut Serum Replacement | Thermo Fisher Scientific | | 10828028 |
| L-Ascorbic acid | Tocris | | 4055 |
| LDN193189 | Sigma | | SML0559-5MG |
| L-glutamine | Thermo Fisher Scientific | | 25030-081 |
| Lyophilized moo glue powder | Modernist Pantry | | N/A |
| Magnesium chloride | Sigma | | M8266-100G |
| Matrigel | BD Biosciences | | 354277 |
| Methyl cellulose,viscosity 15 cPs | [Alfa Aesar](https://my.labguru.com/catalog/companies/1003120) | | 45490-22 |
| N2 Supplement | Thermo Fisher Scientific | | 17502-048 |
| Neurobasal^TM^ medium | Gibco | | 21103-049 |
| Non-essential Amino Acids | Thermo Fisher Scientific | | 11140050 |
| paraformaldehyde | Sigma | | P6148-1KG |
| phosphocreatine | Sigma | | [P7936](https://www.sigmaaldrich.com/US/en/product/sigma/p7936) |
| Potassium chloride | Sigma | | P3911-500G |
| Purmorphamine | Tocris | | 4551 |
| Recombinant human BDNF | Peprotech | | 450-02 |
| Recombinant human GDNF | Peprotech | | 450-10 |
| ROCK inhibitor Y27632 | Stemcell Tech. | | 72304 |
| SB431542 | Stemgent | | 04-0010-10 |
| Shh C25II | R&D Systems | | 464-SH-01M |
| Sodium alginate | Sigma | | W201502 |
| Sodium chloride | Sigma | | S9888-500G |
| Sucrose | Sigma | | S9378-1KG |
| TeSR-E8 medium | Stemcell Tech. | | 05991 |
| Thrombin | Sigma | | T7009 |
| Triton-X100 | Sigma | | X100-500ML |
| UltraPure 0.5 M EDTA | Life Technologies | | 15575-038 |
| VPA | Sigma | | P4543 |
| β-mercaptoethanol | Sigma | | M7522-100ML |
| Critical commercial assays | |  | |
| Lenti-X™ Concentrator | TakaRa Bio | | 631232 |
| *Continued* |  | |  |
| **REAGENT or RESOURCE** | **SOURCE** | | **IDENTIFIER** |
| Lenti-X™ GoStix™ Plus | TakaRa Bio | | 631280 |
| Live/Dead® staining kit | Thermo Fisher Scientific | | R37601 |
| Recombinant DNA |  | |  |
| pAAV.Syn.NES-jRGECO1b.WPRE.SV40 | Addgene | | 100857 |
| pDM2.g | Addgene | | 12259 |
| pENN.AAV.GFAP.iGluSnFr.WPRE.SV40 | Addgene | | 98930 |
| psPax2 | Addgene | | 12260 |
| Experimental models: Cell lines |  | |  |
| Human : H9 ES cells | WiCell Research Institute | | WA09 |
| Human : Syn-GFP-H9 ES cells | WiCell Research Institute | | [[NIHhESC-10-0062](http://grants.nih.gov/stem_cells/registry/current.htm?id=414)](https://www.wicell.org/home/stem-cells/catalog-of-stem-cell-lines/wa09.cmsx) |
| Human : WIZ04e-CAG-mCherry-H9 ES cells | WiCell Research Institute | | WIZ04e |
| Human : CAG-ChR2-YFP-H9 ES cells | WiCell Research Institute | | [WAe009-A-29](https://hpscreg.eu/cell-line/WAe009-A-29) |
| Human : AxD R88C iPS cells | Waisman center iPSC core | | WC-01-01-AL-AM |
| Human: HEK-293T | CCHMC Pluripotent Stem cell core /  ATCC | | CRL-3216 |
| Software |  | |  |
| CorelDraw | Alludo | | N/A |
| Excel | Microsoft | | 2013 |
| Igor | WaveMetrics | | v4.0 |
| ImageJ&FIJI | NIH | | N/A |
| Imaris | Oxford Instruments | | v10.1 |
| MiniAnalysis | WaveMetrics | | v6.0 |
| Notepad++7 | Microsoft | | v7 |
| pClamp | Molecular Devices | | v11.0.3 |
| Prism | GraphPad | | v5&v8 |
| Repetier Host | Hot-World GmbH & Co. KG | | v2.0.5 |
| Slic3r | N/A | | v3 |
| Others |  | |  |
| 6-Well tissue culture plates | Thermo Fisher Scientific | | 140675 |
| 24-Well tissue culture plates | Thermo Fisher Scientific | | 142475 |
| 27G Micronozzle | Cellink | | NZ3270005001 |
| 30G Microneedle | Cellink | | NZ5300505001 |
| 3 mL Cartridge | Cellink | | CSC010311101 |
